# Summer and autumn photosynthetic activity in High Arctic biological soil crusts and their winter recovery

**DOI:** 10.1101/2025.09.09.675080

**Authors:** E. Hejduková, E. Pushkareva, J. Kvíderová, B. Becker, J. Elster

## Abstract

**Introduction:** Biological soil crusts, found in arid and semi-arid areas worldwide, play a crucial role in the carbon cycle. This study analysed biocrusts from three different altitudes in Svalbard (High Arctic) in 2022–2024.

**Methods and Results:** Monitoring of microclimatic parameters, including irradiance, humidity, air, and soil temperature revealed unexpected extremes at the lowest elevation site. Molecular methods were used to determine the diversity of microalgae, revealing the presence of Trebouxiophyceae and Chlorophyceae as the dominant eukaryotic algal groups. Among the cyanobacteria, the dominant taxonomical groups were Nostocales, Pseudanabaenales, and Oscillatoriales. Measured photosynthetic activity was largely driven by irradiance across the different seasons and locations. Higher maximum quantum yield (F_V_/F_M_) values (approximately 0.6) were measured at lower irradiance levels (<100 µmol m^−2^ s^−1^). Photosynthetic activity was observed in early October 2022, and diurnal changes were even noticeable at subzero temperatures in late October 2023, with the low irradiance curve being mirrored by the development of F_V_/F_M_. Furthermore, thawed biocrusts in winter exhibited the ability to rapidly restore photosynthetic activity, which was also supported by the expression of photosynthesis-related genes. Metatranscriptomic analysis revealed that the differential gene expression observed for the D1, RbcS, Ohp1, and ELIP proteins suggests that light stress induced photoinhibition plays a major role in biocrusts, particularly in winter.

**Conclusion:** The biocrusts can remain active for extended periods and provide carbon fixation during times when tundra plants primarily engage in respiration, making them very important for the polar environment.

## 1 Introduction

Biological soil crusts (biocrusts) are communities of microscopic (cyanobacteria, algae, fungi, bacteria) and macroscopic (lichens, mosses, liverworts) organisms living on or within the uppermost millimetres of the soil surface forming a compact layer. Biocrusts are important for maintaining the health and resilience of terrestrial ecosystems as they stabilise soil, enhance soil fertility, and influence local hydrological cycles (Evans and Johansen 1999; Bowker *et al*. 2008; Lichner *et al*. 2013; Bu *et al*. 2014; Williams *et al*. 2017; Gharemahmudli *et al*. 2024). Despite the importance of biocrust and soil microalgae, their field of study is relatively recent and is currently experiencing a surge in interest (Joseph and Ray 2024). However, studies in the polar regions remain limited, and our understanding of the seasonal and diurnal photosynthetic activity of biocrusts incomplete.

In the polar regions, including both the Arctic and Antarctic, biocrusts create patchy or continuous cover that are dominated by bryophytes, lichens, eukaryotic algae and prokaryotic cyanobacteria (Pushkareva *et al*. 2015; Williams *et al*. 2017; Weber *et al*. 2022) and a large number of species have been identified using molecular techniques such as metabarcoding and metagenomics (Rippin *et al*. 2018a; Pushkareva *et al*. 2022, 2023). Specifically, High Arctic polar desert crusts are often dominated by eukaryotic algae and cyanobacteria (Belnap and Lange 2003; Pushkareva *et al*. 2015, 2017, 2023). Naturally, the species composition of biocrusts in polar environments changes during different succession stages of soil development and/or due to the type and chemical composition of the substrate (Pushkareva *et al*. 2015, 2022, 2023; Pessi *et al*. 2019). In the context of global warming and climate change, it is likely that temperatures will increase significantly (Huntington *et al*. 2005; Meredith *et al*. 2019), which may boost microbial activity and diversity. However, extreme conditions such as heatwaves, rain-on-snow events, drought, or flooding can disrupt these communities (Aransiola *et al*. 2024; Bååth and Kritzberg 2024). This could potentially alter the composition of biocrusts, as a previous study of a warm desert suggested, which led to a decrease in the abundance of cyanobacteria (Steven *et al*. 2015).

In polar ecosystems, microbial communities face numerous challenges, including limitations in their capacity for photosynthesis, growth, and reproduction. These challenges involve intense solar radiation during summer (including damaging UV radiation), extended periods of prolonged darkness in winter, low nutrient supply, periods of desiccation, and freezing temperatures (Thomas *et al*. 2008; Pichrtová *et al*. 2020). Therefore, microorganisms have developed physiological and molecular adaptations to survive and thrive in such harsh environments (Morgan-Kiss *et al*. 2006; De Maayer *et al*. 2014; Pichrtová *et al*. 2020) For example, polar algae and cyanobacteria are resistant to abiotic stresses such as freezing, desiccation, UV light, and nitrogen starvation (Davey 1989; Hawes *et al*. 1992; Šabacká and Elster 2006; Elster *et al*. 2008; Tashyreva and Elster 2015; Holzinger *et al*. 2018; Hejduková and Nedbalová 2021; Hejduková *et al*. 2024). Remarkably, some of them tolerate extremely low temperatures (–40 °C and lower) or even –196 °C – temperature of liquid nitrogen (Šabacká and Elster 2006; Elster *et al*. 2008; Hejduková *et al*. 2019, 2024).

Ecological studies of polar algae and cyanobacteria mainly focus on their annual development, morphology, and/or survival mechanisms (Pichrtová *et al*. 2016; Tashyreva and Elster 2016; Hejduková *et al*. 2020), but have not investigated photosynthetic performance in detail. The photosynthetic activity of algae and cyanobacteria from polar and alpine biocrusts has been rarely studied, and mostly under laboratory conditions (Karsten *et al*. 2010; Karsten and Holzinger 2012). As a result, understanding of *in situ* photosynthetic processes in the polar regions is limited to two biocrust studies focussing on the summer growing season in Svalbard (Sehnal *et al*. 2014; Pushkareva *et al*. 2017). In these studies, irradiance appeared to be the main controlling factor of photosynthetic activity, therefore making changes in seasonal and diel dynamics a major environmental parameter. A thorough investigation is necessary to understand the seasonal and diurnal dynamics that have yet to be explored.

This study compares *in situ* diurnal changes in photosynthetic activity of biological soil crusts during summer and autumn 2022–2023 at three localities at different altitudes in Svalbard (High Arctic). Additionally, winter frozen samples collected in March 2023 and 2024 were thawed *“ex situ”* and photosynthetic activity was monitored to explore the recovery from the dormant winter state. We hypothesized that photosynthetic activity reflects the influence of diurnal and seasonal changes in environmental factors such as light availability and temperature. The abundance and diversity of microbial phototrophs including microalgae (term microalgae in the text refers to eukaryotic algae and prokaryotic cyanobacteria, unless further specified), lichenised microalgae and moss development stages and their photosynthesis-related gene transcripts were also evaluated in relation to environmental factors.

## 2 Material and Methods

### 2.1 Study sites description

For this study, three experimental sites were established in the vicinity of Longyearbyen, West Spitsbergen (Svalbard archipelago, High Arctic), in August 2022 (Fig. 1, 2). A description of the studied sites and their microbial community composition is present in Pushkareva *et al*. (2024b). In summary, Site 1 was located in the Bjørndalen valley and two other sites on the slopes of the Breinosa Mountain in the vicinity of Mine 7 (Site 2) and the Kjell Henriksen Observatory (Site 3) (Fig. 1, Supplement S1). Areas with a minimum of 80 % cover of biological soil crust were chosen for the sampling; however, sparse vegetation was also present (Supplement S1). The biocrusts differed macroscopically between the sites (Fig. 2). The biocrusts of the lowest elevated Site 1 were better developed compared to the others with a relatively high diversity of mosses present. It was surrounded by tundra vegetation and the presence of a long-lasting snow cover (snow bed), restricting the growth and distribution of vascular plants. The characteristic of the higher elevated biocrusts at Sites 2 and 3 was different. In the close surroundings, there was only very poor tundra with low cover of vascular plants. Site 2 was represented by a dark, highly compact, and homogeneous crust cover. The biocrust at Site 3 differed from Site 2 by the greater occurrence of lichens. An overview of additional geographic, vegetation, and geological characteristics (Major and Nagy 1972; Dallmann *et al*. 2001; Piepjohn *et al*. 2012) is presented in Supplement S1.

**Fig. 1.**
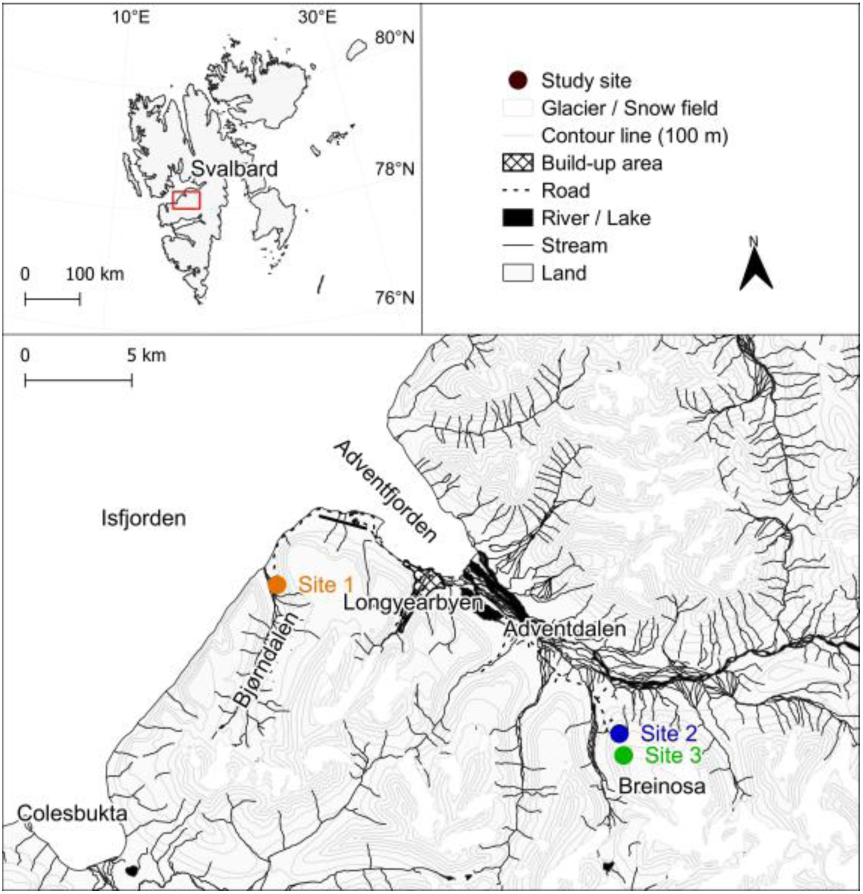
The Svalbard region, with the area of West Spitsbergen (Svalbard archipelago) highlighted. A detailed map illustrating the study sites located near Longyearbyen. Map source: Kartdata Svalbard (Norwegian Polar Institute 2014).

**Fig. 2.**
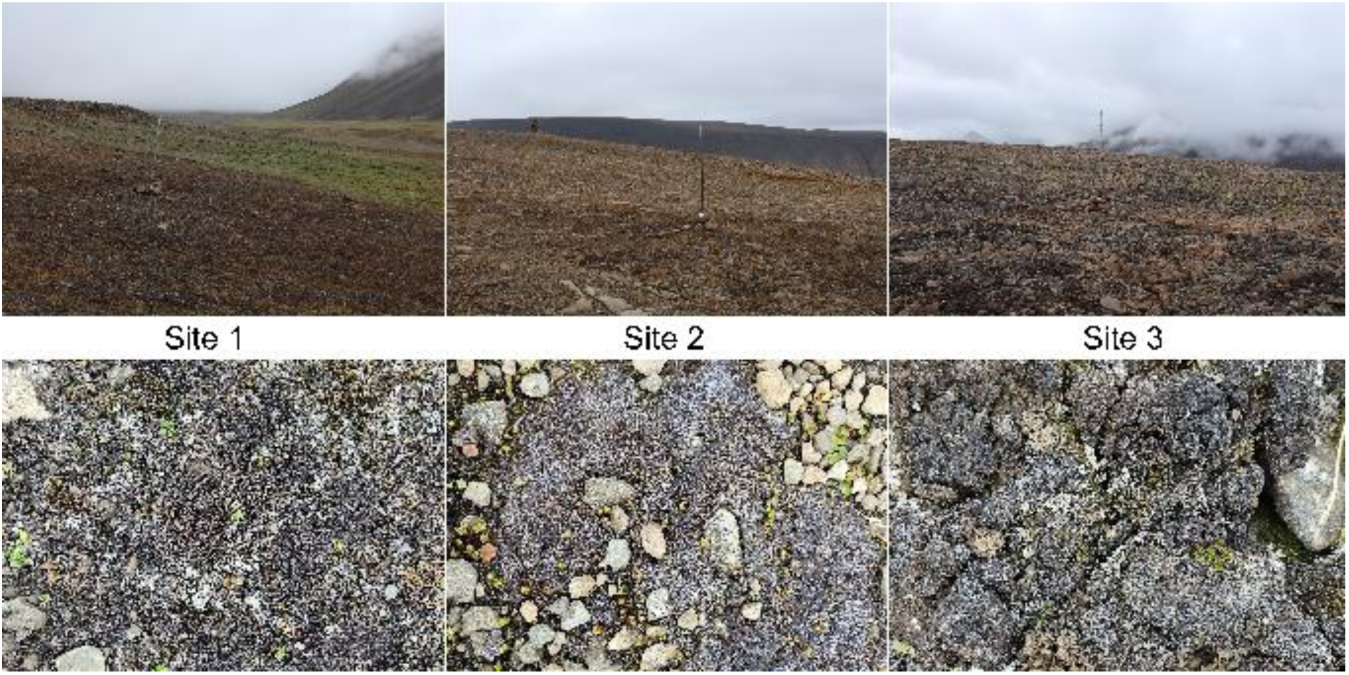
Photographs of the studied sites studied with detailed images of the biological soil crusts.

### 2.2 Microclimate data collection

The climate of West Spitsbergen is classified according to the Köppen-Geiger climate system as semi-arid polar tundra (Førland *et al*. 1997; Peel *et al*. 2007). The Norwegian Meteorological Institute (Norwegian Meteorological Institute 2023) reported that in the period 2010–2020, the average annual precipitation (at the Svalbard Airport meteorological station) was 221 mm, with maxima in August and minima in May. According to the temperatures measured 2017–2022 at Svalbard Airport (Norwegian Meteorological Institute 2023) and in Advent Valley (Adventdalen) provided by the University Centre in Svalbard (UNIS weather stations 2023), the coldest month is March with an average temperature of −13 °C (average minima of −26 °C) and warmest is July with an average temperature of 8 °C (average maxima 15 °C). The mean air temperature exceeds 0 °C for about four months, from the beginning of June until the end of September. Daylight is not available from the end of October to the middle of February.

To provide information on the environment, a series of basic parameters were measured at each site at one-hour intervals throughout the study period. Minikin Tie and later QTHi dataloggers (Environmental Measuring Systems, Brno, Czech Republic) were employed to monitor temperature, photosynthetically active radiation (PAR, in range of 400–700 nm), and air humidity at a height of 120–160 cm above ground. The MicroLog T3 dataloggers were used in conjunction with three soil temperature sensors Pt1000/8 (Environmental Measuring Systems, Brno, Czech Republic) to record the soil temperature at a depth of 2–5 cm below ground. Due to logistical reasons, the PAR and humidity dataloggers were permanently installed later, with Site 1 and 2 in March 2023 and Site 3 in June 2023. Therefore, the data of relative humidity at Site 3 for the first year of study were obtained from the Breinosa weather station (UNIS weather stations 2023) situated next to the original site.

### 2.3 Sampling for metagenomics and metatranscriptomic analyses

Biocrust samples, each 2 cm deep to match the thickness of the biological soil crust, were collected using a sterile laboratory spoon on 5,6,8/8/2022, 3/10/2022, 23/3/2023 and 13/8/2023. To preserve RNAs for molecular analyses, 1 g of the biocrust was placed into a cryotube with 2 ml of LifeGuard Soil Preservation Solution (QIAGEN, Germantown, MD, USA). In March 2023, all sites were covered in frozen snow and, thus, only samples from Site 1 could be retrieved. Five replicates were collected for each of the analyses. The samples were kept at – 20 °C and transported frozen in the laboratories of the Institute for Plant Sciences, of the University of Cologne (Germany).

### 2.4 *In situ* photosynthetic activity measurement

*In situ* photosynthetic activity of the biocrusts and its variation over the day was studied both in summer and autumn in 2022 and 2023. To measure the same area, the biocrusts of the localities were carefully transferred in 15 cm diameter Petri dishes and plastic bowls perforated at the bottom in advance and placed back to their original location (Supplement S2). Three to four bowls and dishes were randomly established within the area of each site as replicates. For comparison between sites only Petri dishes data were used and for evaluation of Site 1 both bowls and dishes were included in analyses, unless further specified.

In the summer seasons, measurements were performed at the three study sites for 24h in 6h intervals at each site on 9–10/8/2022 and 5–6/8/2023. In autumn 4/10/2022 and 23/10/2023 for 7h and 4.5h in intervals of 4h and 1.5h, respectively. Only Site 1 was measured for photosynthetic activity in the autumn period, as Site 2 and Site 3 could not be measured due to frozen snow cover at the higher elevations.

In the field, photosynthetic activity was measured at eight random spots per dish/bowl (in total 32–48 spots per site) using a hand-held FluorPen FP-100 fluorometer (Photon Systems Instruments, Drásov, Czech Republic). The photosynthetic activity was measured using the OJIP protocol after 15 min of dark acclimation. The maximum quantum yield of the photosystem II (φ_Po_ ∼ F_V_/FM) was determined according to Strasser *et al*. 2000a, 2004:

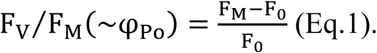

Where F_0_ is the minimum fluorescence at the beginning of OJIP measurement and F_M_ is the maximum reached fluorescence during OJIP transient.

Moreover, the OJIP protocol evaluates various parameters including the following measured and calculated parameters used for the further data analyses on photosynthesis performance: M_0_, V_I_, V_J_, ψ_ET2o_, φ_ET2o_, φ_Do_, J_0_^ABS^/RC, J_0_^TR^/RC, J_0_^ET2^/RC, J_0_^DI^/RC. Their physiological meanings adopted from Stirbet *et al*. 1998; Strasser *et al*. 2004 are listed in Supplement S3.

Additionally, the maximum possible relative electron transport rate (rETR_max_), which is a raw proxy of the maximum capacity of photosynthetic activity, was calculated using computed values of F_V_/F_M_, actual irradiance (PAR), and a factor of 0.5 reflecting the partitioning of energy between photosystems (Maxwell and Johnson 2000) as:

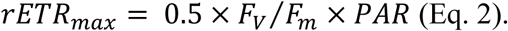

### 2.5 *Ex situ* recovery of photosynthetic activity after winter biocrusts thawing

Furthermore, *ex situ* photosynthetic activity of the thawed biocrusts was measured in the winter season at the end of March in 2023 (23–31/3/2023) and 2024 (25–31/3/2023). Four additional bowls were established at Site 1 as previously described. The biocrusts were extracted from the frozen soil, protected from light and allowed to thaw slowly at a temperature of 4 °C for a period of approximately 24–36 hours. Once the spots free of ice and snow had emerged, the crust were placed at 15 °C and 26 µmol m^−2^ s^−1^ and the effective quantum yield of photosystem II (Ф_PSII_) was carried out at 5-minute intervals on three spots by Monitoring Pen MP-100 fluorometer (Photon Systems Instruments, Drásov, Czech Republic) per bowl/biocrust (in total 12 spots) at the same time for the period of one hour when stable Ф_PSII_ was reached. The Ф_PSII_ was calculated as:

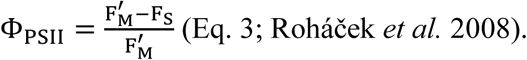

Where F_S_ is the steady-state fluorescence in light and F^′^ is the maximum fluorescence after saturation pulse in light.

After the measurements, the biocrusts were returned to their original location and covered in snow. The identical cycle of thawing and measurement of the biocrusts was repeated after four days.

### 2.6 Metagenomic and metatranscriptomic analyses

The molecular analyses were performed as described in Pushkareva *et al*. 2024b. In summary, DNA extraction was performed using the DNeasy PowerSoil Pro Kit (QIAGEN, Germantown, MD, USA) according to the manufacturer’s instructions. RNA extraction was performed using the NucleoBond RNA Soil Mini kit (Macherey-Nagel Germany). Metagenomic and metatranscriptomic sequencing was conducted at the Cologne Centre for Genomics (Cologne, Germany) using NovaSeq6000 sequencing system (PE150). Of the samples collected in October 2022, only five were sequenced (two replicates from Site 1, one replicate from Site 2, and two replicates from Site 3). Sequences are available in the Sequence Read Archive under the project numbers PRJNA1124630 and PRJNA1172564 for metagenomics and metatranscriptomics, respectively.

### 2.7 Data analyses

Fluorescence data were retrieved by FluorPen 1.1.2.6 software (Photon Systems Instruments, Drásov, Czech Republic). If one or more of the OJIP parameters were out of the range defined in Supplement S3, the computed values were excluded from the analyses. Fluorescence performance was correlated to environmental data and differences among the sites or between seasons were tested using unpaired t-tests or one-way ANOVA with post hoc comparisons using Tukey’s multiple comparison test. PCA was run to summarize the variability within the F_V_/F_M_ values and other non-photochemical parameters of fluorescence: M_0_, V_I_, V_J_, ψ_ET2o_, φ_ET2o_,φ_Do_, J_0_^ABS^/RC, J_0_^TR^/RC, J_0_^ET2^/RC, J_0_^DI^/RC. The effects of site, sampling season, air temperature, soil temperature, and irradiance were tested using RDA with Monte-Carlo permutation test to show statistical significance. Prior to running the PCA and RDA, the data were standardized across species (mean variance standardization).

Bioinformatic analyses were performed in OmicsBox software (v3.3.1) as described in Pushkareva *et al*. 2024b. The rRNAs were separated from both quality-filtered datasets using SortMeRNA (Kopylova *et al*. 2012) and the remaining reads were separately assembled de novo using MEGAHIT (v1.2.8, Li *et al*. 2015). Taxonomic assignment of 16S and 18S rRNAs retrieved from the metagenomic dataset was performed using the Silva database (v138.1) available at SILVAngs analysis platform. The sequences assigned to the cyanobacteria and algae were then retrieved to calculate relative abundance. Moreover, the transcripts and metagenomic contigs were quantified using the RSEM software package (v1.3.3, Li and Dewey 2011) and aligned to the NCBI Blast searches (E × 10). Additionally, Gene Ontology mapping and annotations were performed (Götz *et al*. 2008). Subsequently, the photosynthesis-related genes were retrieved for further analysis.

The influence of environmental parameters on photosynthesis-related genes was tested. A principal component analysis (PCA) was performed to summarise the variability within the relative abundances of photosynthesis-related transcripts. To test the effect of site and sampling season (Aug22 × Oct22 × Mar23 × Aug23) redundancy analysis (RDA) with Monte-Carlo permutation test to show the statistical significance was used. Prior to running the PCA and RDA, the data were standardized across species (mean variance standardization). The impact of site, sampling season and their interaction on photosynthesis-related transcripts represented by the FPKM (fragments per kilobase of transcript per million fragments sequenced) numbers was tested using two-factor analyses of variance (ANOVA).

Statistical analyses were performed using R 4.4.1 (R Core Team, Vienna, Austria), Statistica 14.0 (TIBCO Software, San Ramon, CA, USA) or GraphPad Prism 5.03 (GraphPad Software, La Jolla, CA, USA). The ordination analyses were performed in Canoco 5.01 (Biometris, Wageningen, Netherlands, Ter Braak and Šmilauer 2012). The environmental measurement data were processed with the Mini32 program Mini32 (Environmental Measuring Systems, Brno, Czech Republic). For further visualization of the data the following software was used: QGIS 3.28 (Quantum GIS Geographic Information System, London, UK), GraphPad Prism 5.03 (GraphPad Software, La Jolla, CA, USA), SigmaPlot 14.0 (Grafiti, Palo Alto, CA, USA), Zoner Photo Studio 16 (Zoner Software, Brno, Czech Republic) and Inkscape 1.1 (Software Freedom Conservancy, New York, NY, USA).

## 3 Results

### 3.1 Temperature, relative humidity, and irradiance

The mean daily values of air and soil temperature, relative air humidity, and PAR from 6/8/2023 to 24/10/2023 are presented in Fig. 3 and their statistical significance of differences among the three study sites is shown in Table 1. Overview of microclimate data including month means, minimum and maximum values are presented in Supplement S4 and in Pushkareva *et al*. 2024b.

**Fig. 3.**
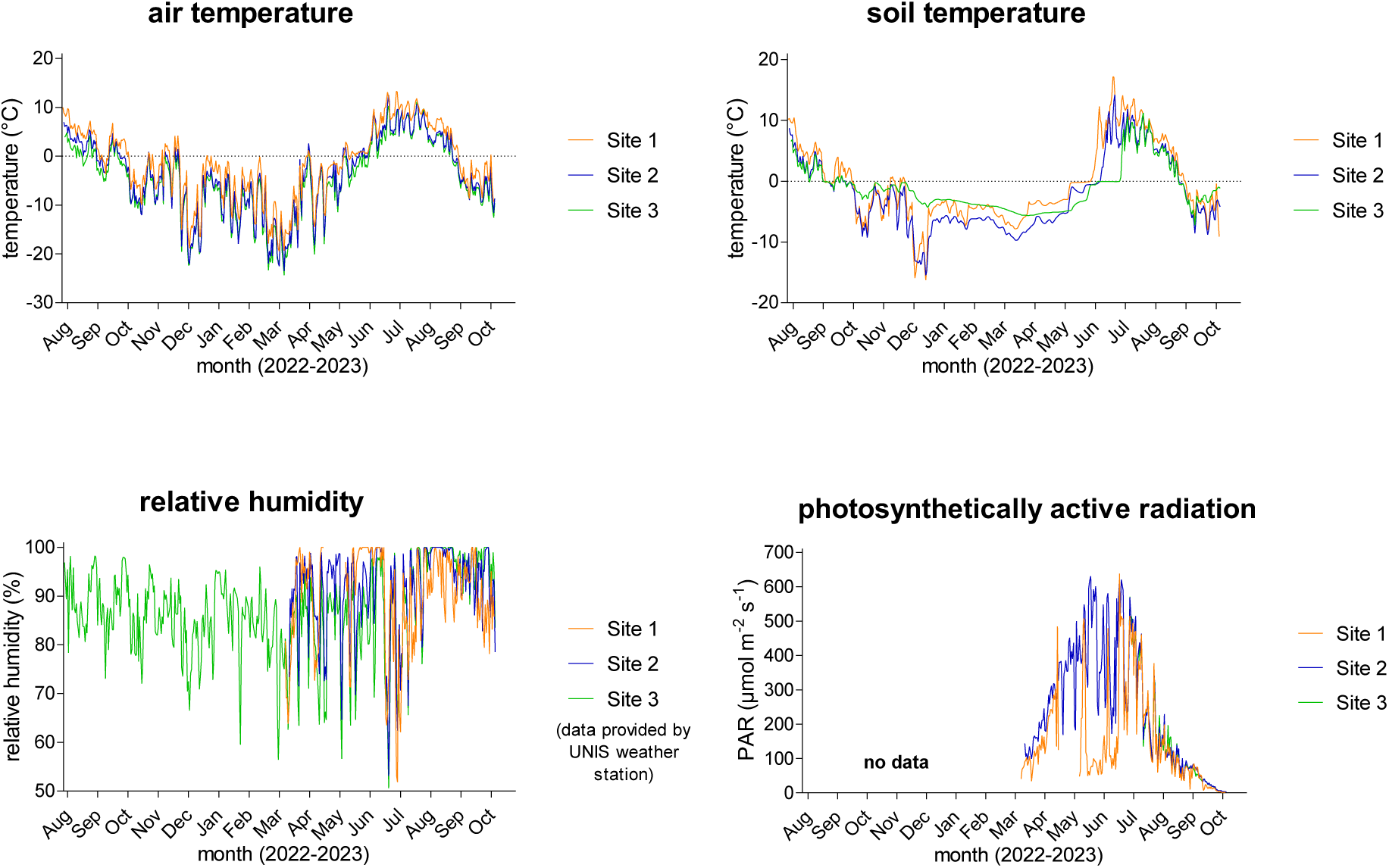
Diel averages of air and soil temperatures, relative humidity, and photosynthetically active radiation (PAR) from 6/8/2023 to 24/10/2023.

**Table 1.** The statistical significance (unpaired t-tests; n = 442) of the parameters monitored between the three study sites from 6/8/2023 to 24/10/2023. The statistically significant differences are marked in bold.

|  | Air temperature | Soil temperature | Relative humidity | Photosynthetically active radiation |
| --- | --- | --- | --- | --- |
| Site 1 × Site 2 | **P < 0.0001, t = 4.809** | **P < 0.0001, t = 4.729** | P = 0.1084, t = 1.609 | **P < 0.0001, t = 5.977** |
| Site 1 × Site 3 | **P < 0.0001, t = 6.804** | **P = 0.0242, t = 2.258** | P = 0.1501, t = 1.442 | P = 0.7558, t = 0.311 |
| Site 2 × Site 3 | P = 0.0538, t = 1.931 | **P = 0.0006, t = 3.453** | **P = 0.0020, t = 3.110** | **P < 0.0001, t = 5.102** |

The daily mean air temperature at Site 1 ranged from −19.5 °C (March 2023) to 13.3 °C (July 2023), at Site 2 from –23.5 °C (March 2023) to 12.5 °C (July 2023), and from –24.4 °C (March 2023) to 11.1 °C (August 2023) at Site 3. The lowest air temperature of −26.9 °C was measured on 21/3/2023 at Site 3 and the air temperature rose to 16.3 °C on 5/7/2023 at Site 1. Regarding the soil temperature at Site 1 the lowest daily mean was –16.2 °C (December 2022) and the highest 17.1 °C (July 2023), at Site 2 ranged from –15.4 °C (December 2022) to 14.2 °C (July 2023) and at Site 3 –7.2 °C (September 2023) to 11.2 °C (July 2023). The lowest soil temperature reached –17.4 °C on 25/12/2022 and the highest 24.5 °C on 5/7/2023, both measured at Site 1, making it surprisingly the most extreme site. The air and soil temperature positively correlated at all three sites: P < 0.0001, r = 0.9060 (Site 1), r = 0.8688 (Site 2), r = 0.7340 (Site 3), n = 442. The air temperature was significantly higher at Site 1 than at Sites 2 and 3 where the air temperature was comparable. On the other hand, all sites differed significantly in soil temperature (Table 1).

During the study period, the mean relative humidity of the air exceeded 80 %. As seen in Fig. 3, daily means varied between 50 % and 100 %. The highest variation of relative humidity and daily minimum values were observed from the beginning of January to the beginning of July. Unfortunately, the air humidity dataloggers were not installed at the very beginning of the field study at Site 1 and 2, therefore some data for Sites 1 and 2 are missing. Nevertheless, the relative humidity differed significantly at Sites 2 and 3, but both sites were comparable to Site 1 (Table 1).

The seasonal variation of photosynthetically active radiation day averages ranged from 0 to about 640 µmol m^−2^ s^−1^, which can be expected from such a geographic location due to changes in the solar elevation angle and the Earth–Sun distance throughout the year (Fig. 3). The highest intensity reached 1699 µmol m^−2^ s^−1^ and was measured on 29/4/2023 at Site 1. Similarly to relative humidity sensors, the irradiance sensors were additionally installed only for the 2023 summer season and unfortunately some issue appeared at Site 1, as extremely low levels of intensity and continuous 100 % relative humidity were measured for the most of May. Surprisingly, PAR was similar at the more geographically distant Sites 1 and 3, while it was significantly different between geographically nearby Sites 2 and 3. The significant difference was also observed between Sites 1 and 2 (Table 1).

### 3.2 Diversity of algae and cyanobacteria

The composition of the microalgal community was determined using the rRNA reads present in the metagenomic data set published by Pushkareva *et al*. 2024b (Fig. 4). Trebouxiophyceae dominated the community of eukaryotic algae (52–69 % of algal reads) (Fig. 4). A total of 22 trebouxiophyte genera were identified, and most reads for this class belonged to *Coccomyxa* (14–27 % of algal reads), *Elliptochloris* (13–17 % of algal reads), and unclassified Trebouxiophyceae (17–21 % of algal reads). Chlorophyceae constituted 13–25 % of algal reads, but most reads could not be identified to the genus level. Furthermore, Xanthophyceae were detected only in Site 2, while Eustigmatophyceae only in Site 3. Furthermore, Site 3 had a more diverse community of Bacillariophyceae (5 identified genera), Chlorophyceae (5 identified genera), Chrysophyceae (7 identified genera), and Trebouxiophyceae (17 identified genera) compared to the other sites. A significant effect of the algal classes was revealed (two-way ANOVA, P < 0.0001, F = 141.4, n = 5), along with a significant group × site interaction (P = 0.0028, F = 1.925, n = 5). The site effect was not significant. The only groups that differed significantly among the sites were Chlorophyceae (one-way ANOVA, P = 0.0464, F = 4.009, n = 5) and Eustigmatophyceae (one-way ANOVA, P = 0.0060, F = 8.067, n = 5).

**Fig. 4.**
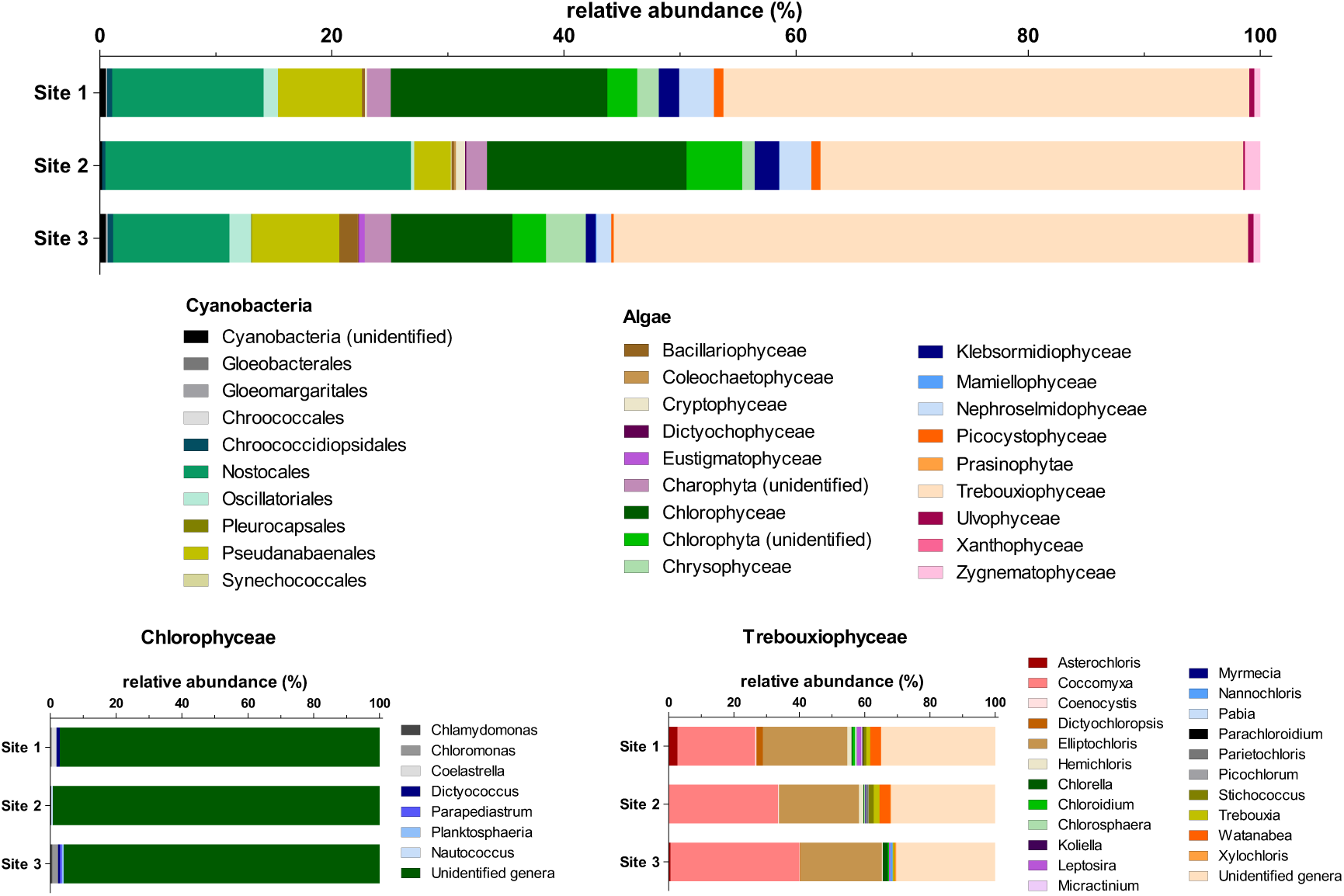
Relative abundances (based on the number of 16S or 18S rRNA reads present, n = 5) of dominant algal classes and cyanobacteria order per study site, with Chlorophycean and Treuboxiophycean genera shown below.

Regarding cyanobacteria, the most dominant groups were Nostocales (49–87 % of cyanobacterial reads) with 12 identified genera and Pseudanabaenales (10–37 % of cyanobacterial reads) involving 13 genera. Gloeobacterales, Gloeomargaritales and Pleurocapsales were identified only on Site 3 and Synechococcales on Site 2. Similarly to algal relative abundances, the cyanobacterial orders significantly differed (two-way ANOVA, P < 0.0001, F = 110.5, n = 5) as well as an interaction between group and site (P < 0.0001, F = 3.773, n = 5). The site effect alone was not significant. Nostocales were the only group that differed significantly among the sites (one-way ANOVA, P = 0.0262, F = 5.008, n = 5).

### 3.3 *In situ* photosynthetic activity

The *in situ* photosynthetic activity, expressed as maximum quantum yield (F_V_/F_M_) and maximum possible relative electron transport rate (rETR_max_), and environmental data were measured in August and October in 2022 and 2023 for one diel cycle (5 “time points” indicating measurement in August 2022 and 2023, 3 “time points” in October 2022 and 4 in October 2023). While the summer measurements were performed at all experimental sites, the autumn measurements were performed only at Site 1 because a deep layer of frozen snow was already present at Sites 2 and 3.

The diel means of F_V_/F_M_ and rETR_max_ in both years were comparable and the differences in biocrust photosynthetic activity among the studied sites in summer were found significant only for F_V_/F_M_ in 2022 (Table 2; one-way ANOVA, P < 0.0001, F = 34.00, n_(Site1)_ = 14, n_(Site2)_ = 19, n_(Site3)_ = 20). Contrary to similarity in photosynthetic activity among the experimental sites, the air temperature in 2022 (Supplement S5; one-way ANOVA, P < 0.0001, F = 98.59, n = 29) and 2023 (Supplement S5; one-way ANOVA, P = 0.0013, F = 7.176, n = 29), soil temperature in 2022 (Supplement S5; one-way ANOVA, P < 0.0001, F = 26.22, n = 29) and the relative humidity in 2023 (Supplement S5; one-way ANOVA, P = 0.0005, F = 8.258, n = 29) significantly differed between the sites during the period of photosynthetic measurements in August.

**Table 2.**
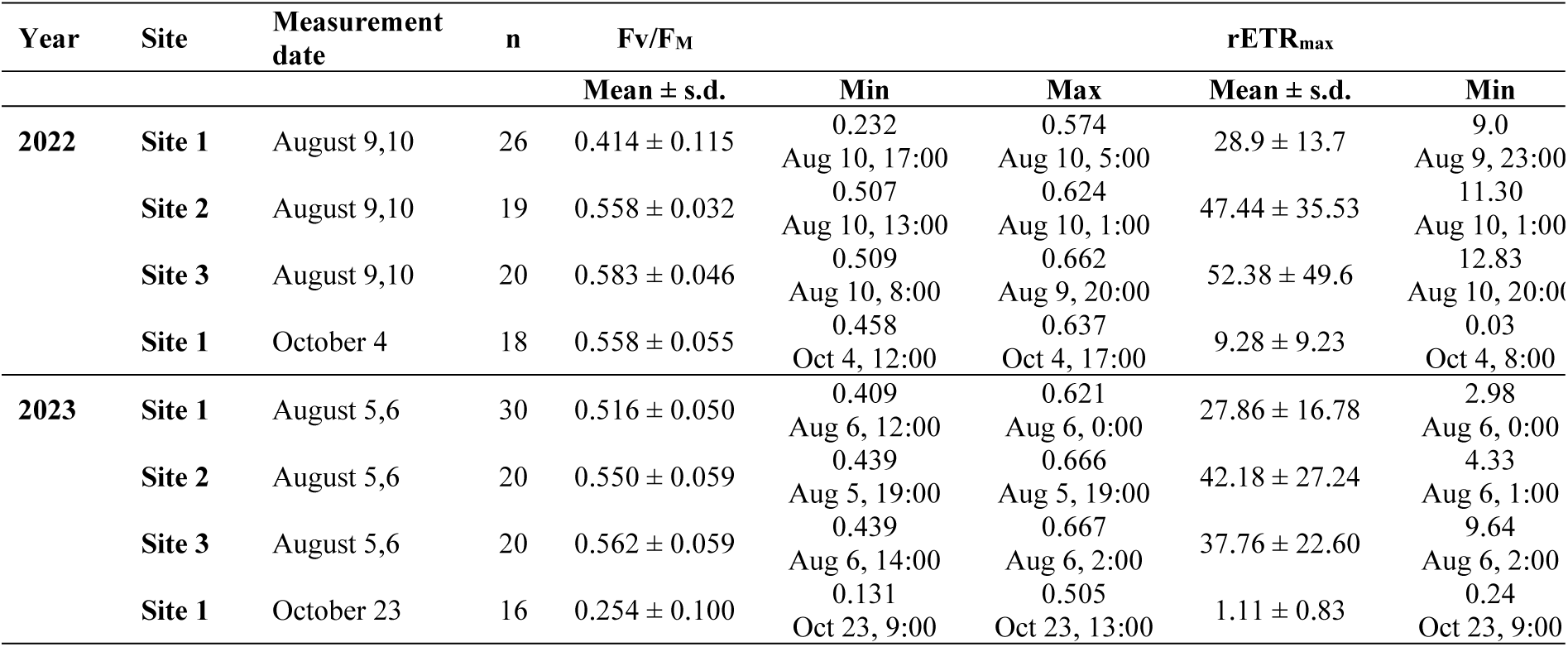
The diel mean (± standard deviation), minimum and maximum values of maximum quantum yield (F_V_/F_M_) and maximum possible relative electron transfer rate (rETR_max_). Date and time (CET) at diel minima and diel maxima indicate when given minimum or maximum was observed. Abbreviation: max – diel maximum, min – diel minimum, n – number of cases. Data used: averages per Petri dish and bowl.

Contrary to summer, the autumn measurements of biological soil crust were performed for two subsequent years (2022 and 2023, both in October) only at Site 1. In general, the values of F_V_/F_M_ measured at the beginning of October 2022 were higher than in the late October 2023 indicating more serious stress encountered in the field in 2023 (Table 2, Supplement S5; unpaired t-tests, P < 0.0001, t = 11.17, n_(Oct22)_ = 18, n_(Oct23)_ *=* 16). Contrary to 2022, the values of rETR_max_ were lower and more stable in 2023 (Table 2, Supplement S5; unpaired t-test, P = 0.0013, t = 3.520, n_(Oct22)_ = 18, n(Oct23) *=* 16). Naturally the measured environmental data also differed significantly: air temperature (unpaired t-test, P < 0.0001, t = 13.77, n = 6), soil temperature (unpaired t-test, P < 0.0001, t = 28.45, n = 6), and irradiance (unpaired t-test, P = 0.0048, t = 3.345, n = 6).

The comparison of the photosynthetic activity of the biocrust among the seasons, performed at Site 1 only, revealed significant differences in both the F_V_/F_M_ (Fig. 5; Kruskal-Wallis ANOVA, P < 0.0001, H = 47.02, n_(Aug22)_ = 26, n_(Oct22) =_ 18, n_(Aug23)_ = 30, n_(Oct23)_ = 16) and the rETR_max_ values (Fig. 5; Kruskal-Wallis ANOVA, P < 0.0001, H = 49.43, n_(Aug22)_ = 26, n_(Oct22) =_ 18, n_(Aug23)_ = 30, n_(Oct23)_ = 16). F_V_/FM differed significantly between all pairs of measurements, with the exception of the beginning of October 2022 and August 2023 when the similar and maximum F_V_/F_M_ values were observed. In August 2022, wide variation in F_V_/F_M_ occurred, while minimum values of F_V_/F_M_ were recorded in October 2023 (Fig. 5). Significant seasonal variation (summer × autumn) in rETR_max_ was observed, with maxima during summer season and minima in autumn. No significance was demonstrated within the two autumn and summer datasets (Fig. 5).

**Fig. 5.**
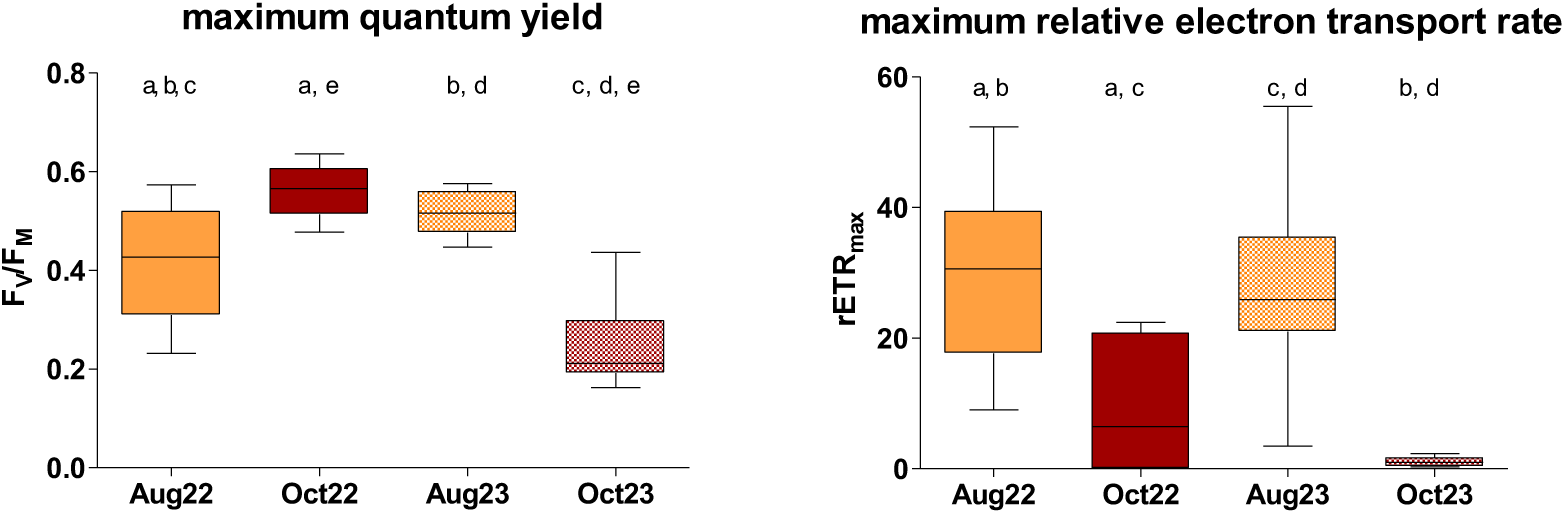
The seasonal comparison of biological soil crust photosynthetic activity (F_V_/F_M_ and rETR_max_) at Site 1. Significant pairs indicated by letters above (n_(Aug22)_ = 26, n_(Oct22) =_ 18, n_(Aug23)_ = 30, n_(Oct23)_ = 16). Line: median; box: first and third quartiles; whiskers: 10–90 percentile.

### 3.4 Diel changes in photosynthetic activity

Despite of continuous light during the polar summer, profound changes in PAR occurred leading to observation of diel cycles at Site 1. Most of the diel changes in “time point” mean values for F_V_/F_M_ ranging from 0.273 to 0.525 and from 0.486 to 0.566 in August 2022 and August 2023, respectively, were statistically significant (Fig. 6, Supplement S5), and these values were comparable between the years. Similarly, the significant changes in mean value for rETR_max_ were observed in range 9.43 to 49.70 and 3.47 to 53.56 in August 2022 and August 2023, respectively, (Fig. 6, Supplement S5), again being comparable between the years. Although the maximum mean values of F_V_/F_M_ were reached around local midnight and early in the morning, the maximum hours mean value for rETR_max_ occurred around local midday in both summer seasons (Fig. 6, Supplement S5).

**Fig. 6.**
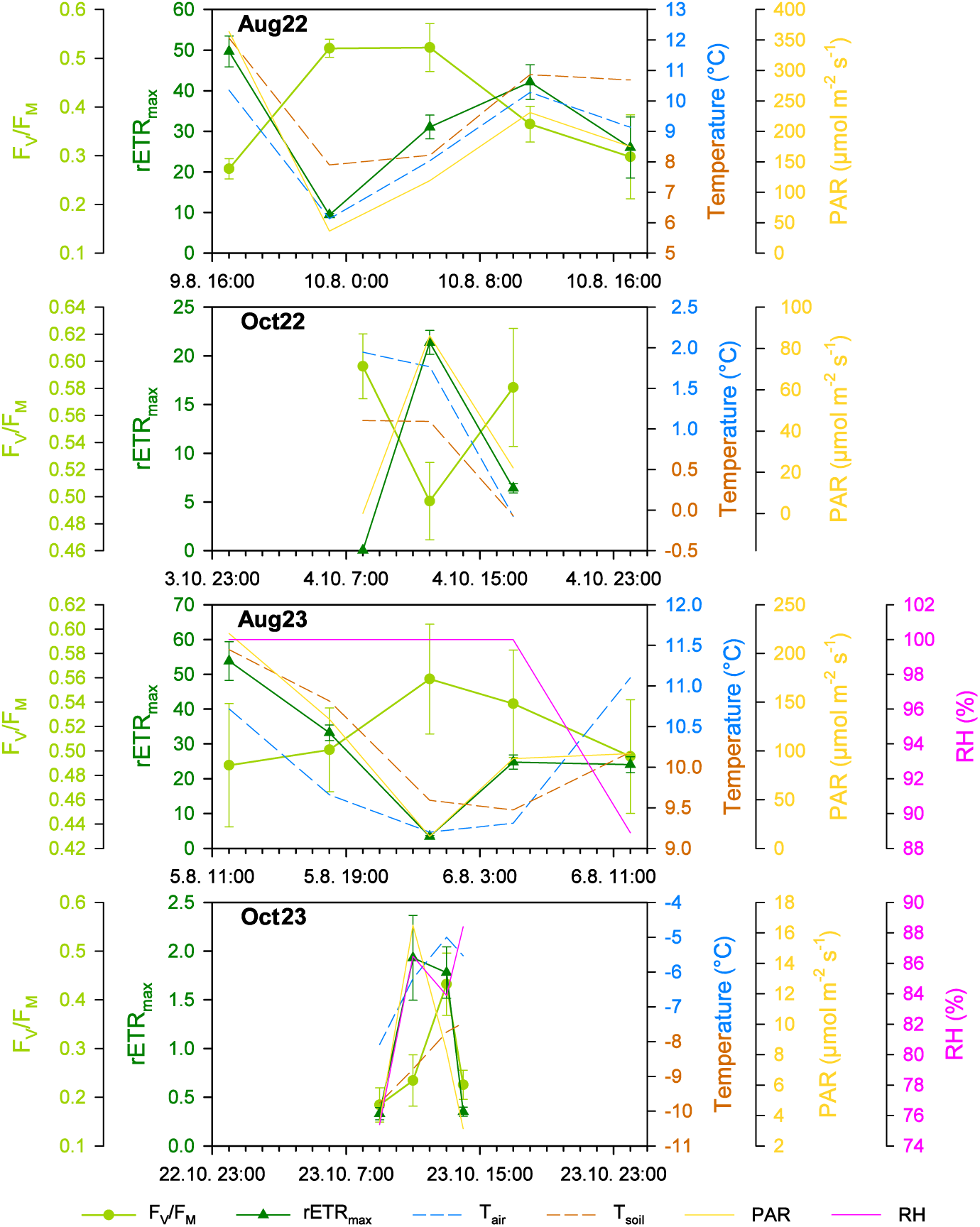
The diel changes of the photosynthetic (F_V_/F_M_ and rETR_max_; mean ± s.d., for n refer to Supplement S5) and environmental parameters (air and soil temperature, T_air_, Tsoil; photosynthetically active radiation, PAR; relative humidity, RH) at Site 1 in the studied periods in summer (Aug) and autumn (Oct) 2022 (22) and 2023 (23).

In autumn, the light period lasted 9 hrs 40 mins on October 4, 2022, and only 3 hrs 34 mins on October 23, 2023. The “time point” mean F_V_/F_M_ was significantly higher in 2022 indicating more stressing conditions in 2023 (Fig. 6, Supplement S5; unpaired t-test, P = 0.0100, t = 4.036, n_(Oct22)_ = 3, n_(Oct23)_ = 4). Despite of low irradiances, especially in 2023, the diel cycles of photosynthetic activity were observed. The mean F_V_/F_M_ in range of 0.496 to 0.597 in early October 2022 were even higher than in summer (Fig. 6, Supplement S5), while in late October 2023, the mean F_V_/F_M_ dropped much lower, to 0.188–0.436 (Fig. 6, Supplement S5). The mean rETRmax of 0.03–21.35 in early October 2022 was reduced to ca 1/3 in comparison considering the summer values (Fig. 6, Table 2, Supplement S5) and in late October 2023, the photosynthetic activity was almost negligible with mean rETR_max_ between 0.34 and 1.99 (Fig. 6, Supplement S5). In early October 2022, the diel course of “time point” mean values of F_V_/F_M_ and rETR_max_ and its response to PAR changes were similar to summer (Fig. 6, Supplement S5), while in late October 2023, the mean F_V_/F_M_ remained low in the local morning while rETR_max_ reached its maximum. At local midday, the higher F_V_/F_M_ and rETR_max_ means were observed than in the local early morning and in the local evening (Fig. 6, Supplement S5).

The diel cycles measured at Sites 2 and 3 in summer revealed similar response as at Site 1, *i*.*e*. maximum “time point” mean F_V_/F_M_ around local midnight and maximum “time point” mean rETR_max_ around local midday (Supplements S5 and S6). At Site 2, the mean F_V_/F_M_ remained stable during the day, spanning from 0.545 to 0.605 and from 0.524 to 0.607 in August 2022 and August 2023, respectively, being slightly higher than at Site 1 (Supplements S5 and S6; unpaired t-test, P = 0.0085, t = 2.956, n = 10) and comparable to Site 3. The “time point” mean rETR_max_ reached its maximum around local midday likewise at Sites 1 and 3 (2022: 11.77– 97.96 and 2023: 4.85–71.35; Supplements S5 and S6). While the mean rETR_max_ minima at Site 2 were comparable to Site 1, the maximum values were higher than at Site 1 and comparable or only slightly lower to Site 3 (Supplements S5 and S6). Likewise, the ”time point” mean F_V_/F_M_ values at Site 3 of 0.529–0.628 and 0.470–0.631 in August 2022 and August 2023, respectively, were higher than at Site 1 at the comparable “time point” (Supplements S5 and S6; unpaired t-test, P = 0.0074, t = 3.015, n = 10) as well as the mean rETR_max_ in ranges of 13.55–139.51 and 10.22–70.67 in August 2022 and August 2023, respectively, however not significant (Supplements S5 and S6).

### 3.5 Effects of environmental parameters on photosynthetic activity

No statistically significant correlations were found among the air or soil temperature, relative humidity and photosynthetic activity expressed as the F_V_/F_M_ and rETR_max_ except for positive correlations of air temperature and rETR_max_ at Site 1 and 3 in August 2022, and negative correlation of soil temperature and F_V_/F_M_ at Site 1 in August 2022 and 2023 (Supplement S7). If significant, negative correlations of F_V_/F_M_, and irradiance were found. Strong significant positive correlations of rETR_max_ and irradiances were found. Interestingly the only significant relationship for autumn measurements was found for rETR_max_ and irradiances in October 2022.

### 3.6 OJIP fluorescence parameters

At all the three sites, positive correlations were found between soil and air temperature, stronger at Sites 1 and Site 3 than at Site 2. None or weak positive correlations of PAR to temperatures were found (Fig. 7).

**Fig. 7.**
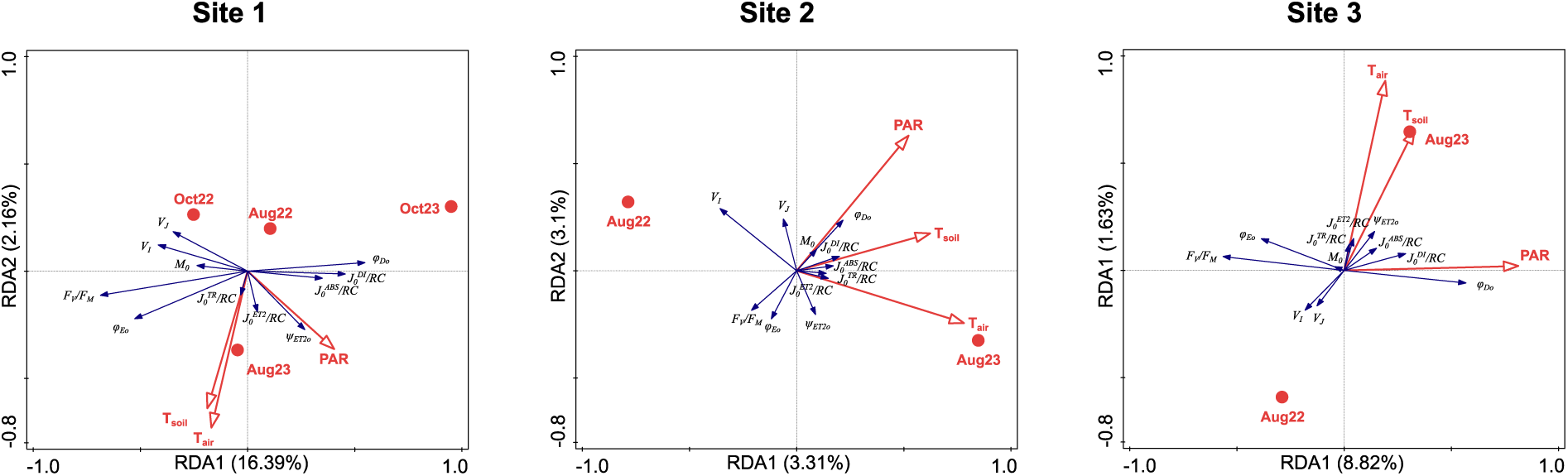
RDA analyses showing the correlation among photosynthetic parameters (explained variables: F_V_/F_M_, M_0_, V_I_, V_J_, ψ_ET2o_, φ_ET2o_, φ_Do_, J_0_^ABS^/RC, J_0_^TR^/RC, J_0_^ET2^/RC, J_0_^DI^/RC; blue arrows) and the environmental data (explaining variables: sampling season, PAR, air and soil temperature; red symbols and arrows) for all the studies sites. Total variation 5247 (Site 1) / 2398 (Site 2) / 3080 (Site 3), explanatory variables account for 19.26 % / 6.79 % / 11.19 %. Monte-Carlo permutation test results: P = 0.002 / P = 0.006 / P = 0.002, pseudo-F = 15.4 / 1.8 / 6.7 (first axis); P = 0.002 / P = 0.02 / P = 0.002, pseudo-F = 18.7 / 3.9 / 8.7 (all axes).

At Site 1, the temperatures and PAR increased to measurement in August 2023. The responses of OJIP parameters differed from Site 2 and 3, as the October measurements were included in the RDA analysis. The J_0_^TR^/RC, J_0_^ET2^/RC and ψ_ET2o_ were positively related to ait and soil temperatures and PAR. The parameters indicating increased energy dissipation, φ_DIo_ and J_0_^DI^/RC, and J_0_^ABS^/RC were related to harsh conditions in October 2023. Opposite reaction was observed in photochemistry related parameters F_V_/F_M_, M_0_ and φ_ET2o_. The parameters V_J_ and VI, were positively related to October 2023. The seasons (Aug × Oct) and years appeared to contribute significantly to data variation, since the explained variation was much higher at Site 1 (Fig. 7).

At Site 2, the air and soil temperatures were increased in August 2023, and the PAR remained independent on the year of measurement and was rather positively correlated to soil temperature. The ψ_ET2o_ and all fluxes through active reaction centre, J_0_^ABS^/RC, J_0_TR/RC, J_0_^ET2^/RC and J_0_^DI^/RC were strongly positively related to air and soil temperatures, while M_0_ and φ_DIo_ were strongly positively related to PAR, weaker positive correlation was found between VJ and PAR. The F_V_/F_M_ and φ_ET2o_ were negatively related to PAR. The V_I_ was strongly negatively related to T_air_ while it was independent on PAR and inclined to measurement in August 2022. (Fig. 7).

Like at Site 2, the T_air_ and T_soil_ raised toward August 2023 at Site 3, but they were more positively correlated. Contrary to Site 2, the relation of PAR was almost independent on both temperatures and on the year of the measurements, but it was slightly positively related to August 2023 measurement. The J_0_^TR^/RC, J_0_^ET2^/RC and ψ_ET2o_ were strongly positively related to increased temperatures in August 2023, while V_J_ and V_I_ were related negatively and inclined towards August 2022. The J_0_^ABS^/RC was positively correlated with temperatures and PAR. The parameters related to energy dissipation, φ_DIo_ and J_0_^DI^/RC, and J_0_^ABS^/RC, increased with higher PAR, opposite reaction was observed in photochemistry related parameters, F_V_/F_M_, M_0_ and φ_ET2o_ (Fig. 7).

### 3.7 *Ex situ* recovery of photosynthetic activity after winter biocrusts thawing

*Ex situ* recovery of photosynthetic activity after winter biocrusts thawing measurements were performed on samples collected at Site 1 at the end of March 2023 and 2024. In both seasons, the effective quantum yields (Φ_PSII_) exhibited a gradual increase during the period of the first hour following both thawing cycles with rapid increase phase within the first 20–35 mins of the exposition and were followed by steady-state later on (Fig. 8, Supplement S8). The maximum measured Ф_PSII_ value reached 0.68 in the first cycle in 2023, while the lowest value was 0.01 in 2024. A notable divergence between the two thawing cycles runs was observed in both years (unpaired t-test; 2023: P = 0.0120, t = 2.718, n = 12, 2024: P < 0.0001, t = 8.792, n = 12). A significant difference was observed between the two years only with regard to the development of the first thawing (unpaired t-test; P < 0.0001, t = 12.70, n *=* 12).

**Fig. 8.**
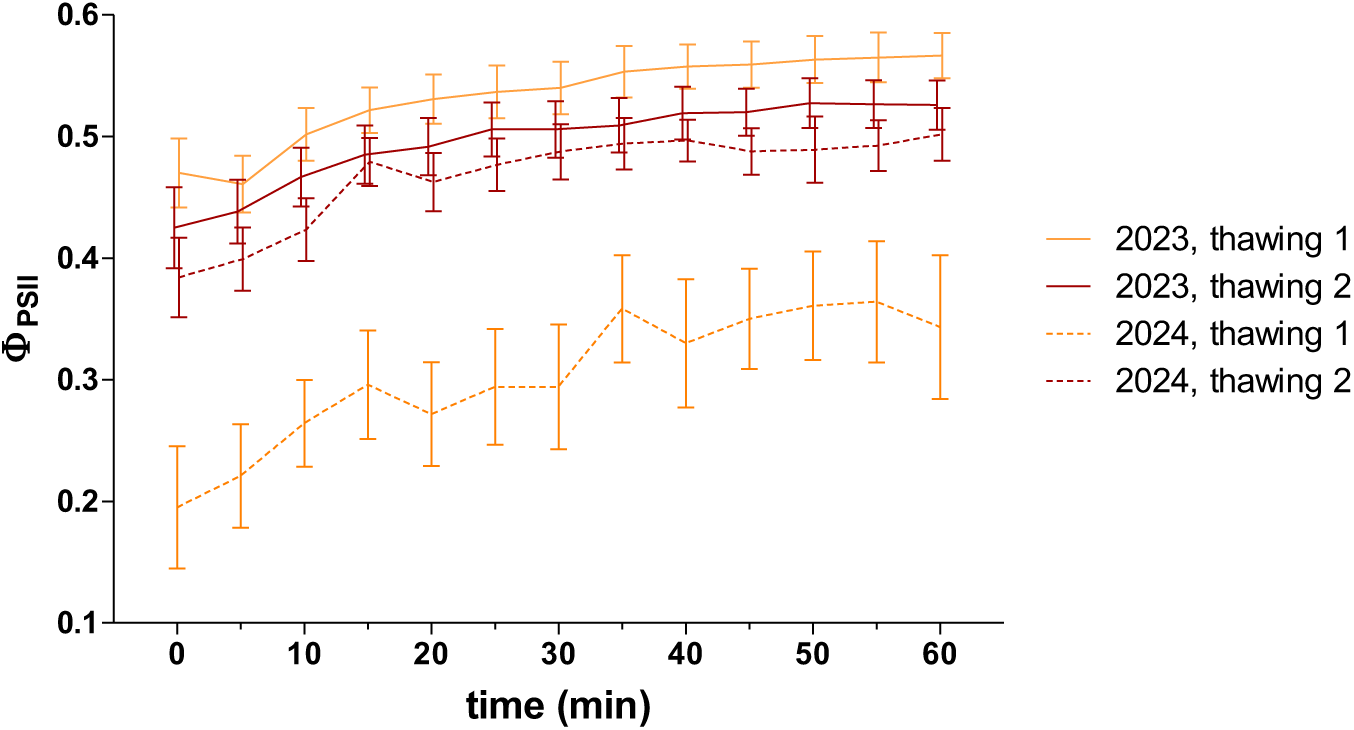
Effective quantum yield of photosystem II (Ф_PSII_; mean ± SEM, n *=* 12) of biocrusts at Site 1 immediately after the thawing cycles in winter (March) 2023 and 2024.

### 3.8 Expression of photosynthesis-related genes

Metatranscriptomic analyses were used to test whether biocrusts acclimate to the different seasons or to the environment of the three sites by differential gene expression. In this study, photosystem II (PSII) related genes and some other photosynthesis or stress-related genes such as oxygen evolving enhancer proteins (OEEs: PsbO, PsbP, PsbQ), RuBisCO small subunit (RbcS), Early Light-Induced Proteins (ELIPs), Cor413PM1 and Ohp1 were studied (Supplement S9) at all three experimental sites in August 2022 and 2023, and October 2022. In March 2023, the samples were collected at Site 1 only.

The *PsbA* (D1 protein coding) and RuBisCo small subunit (*RbcS*) were the most abundant transcripts at the three sites and in all seasons, followed by the other three PSII core subunits *PsbD* (D2 protein coding) and *PsbC* and *PsbB*. All other transcripts showed a much lower relative abundance (Fig. 9). The PSII core subunits (*PsbA*, *PsbB*, *PsbC*, and *PsbD*), four of minor PSII subunits (*PsbH*, *PsbL*, *PsbT*, *PsbZ*), three of the OEEs (*PsbO*, *PsbP* and *PsbQ*), two of the light harvesting complex (*LhcA* and *LhcB*), two involved in PSII assembly (*Ycf48* and *Ohp1*) and *Elip* showed differential gene expression at different sites and part of the year as well as *RbcS*. The expression level of *RbcS* decreased dramatically in March, while the relative abundance of the *PsbA* transcript was maximal in this season (Fig. 9). In particular, the *PsbA* expression was also higher for all three sites in August 2023 compared to August 2022 (Fig. 9).

**Fig. 9.**
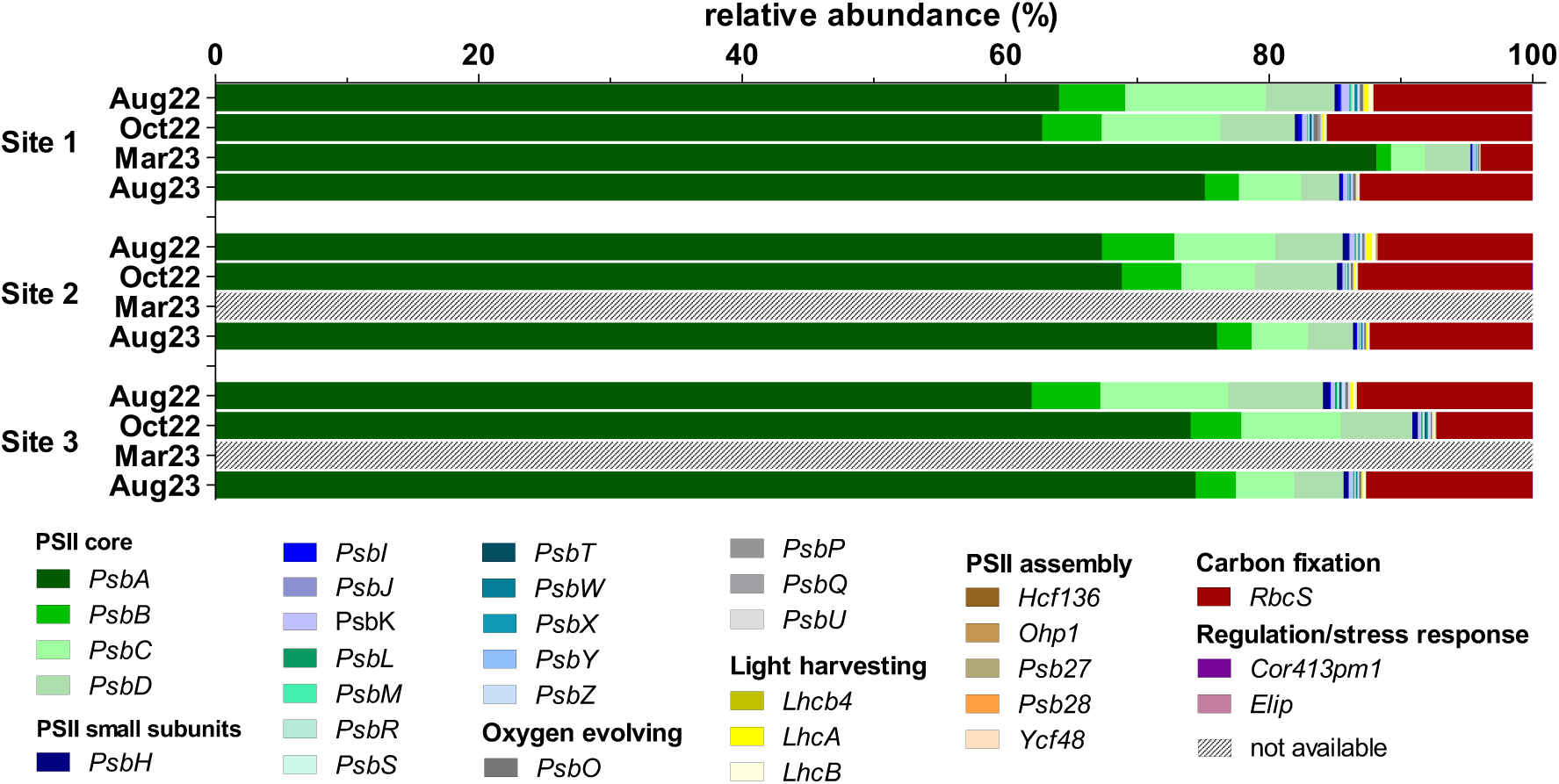
Relative abundance of photosynthesis-related transcripts (expressed in percentage of FPKM, fragments per kilobase of transcript per million fragments sequenced) per study site and sampling season. n_(Aug22)_ = 5, n_(Oct22)_ = 2, n_(Mar23, Aug23)_ = 4.

Of the 32 analysed genes, the expression of 25 transcripts was significantly affected by season and/or the sites and their interactions (Fig. 9, Supplement S10). The effects of site only were significant only in expression of PSII small subunit gene *PsbT* and PSII assembly involved gene *Ycf48*. Gene expression of more transcripts was affected only by season, namely PSII small subunit genes *PsbI*, *PsbJ*, *PsbK*, *PsbM*, *PsbR*, *PsbS*, *PsbW* and *PsbY*, and oxygen evolving complex gene *PsbO*. The effects of season and site were significant in all PSII core genes, in PSII small subunit genes *PsbH* and *PsbZ*, light harvesting complex genes *LhcA* and *LhcB*, and *RbcS*. The season and interaction site × season were significant for the oxygen evolving complex gene *PsbQ* expression. Finally, site, season, and the site × season interaction were important for expression of the PSII small subunit gene *PsbL*, the oxygen evolving complex gene *PsbP* and the PSII assembly-involved gene *Ohp1*. On the contrary, no significant differences in gene expression were observed in the PSII small subunit gene *PsbX*, the oxygen evolving complex genes *PsbU*, the light harvesting complex gene *Lhcb4*, PSII assembly-involved genes *Psb27*, *Psb28* and *Hcf136*, and the desiccation marker *Cor413pm1* (Supplement S10).

The sampling season explained significant part of the variation, 31.31 % (RDA, Monte Carlo permutation test; first axis pseudo-F = 3.2, P = 0.002; all axes pseudo-F = 4.7, P = 0.002; n *=* 35). The effect of site was weaker and explained 11.47 % of variation (RDA, Monte Carlo permutation test; first axis pseudo-F = 1.5, P = 0.006; all axes pseudo-F = 2.1, P = 0.006; n *=* 35). When individual sampling sites were considered, the RDA revealed distinct gene expression during individual season at each site (Supplement S11).

Nevertheless, some general trends in genes expression occurred at all experimental sites. The opposite increase of expression of PSII core genes *PsbA* × *PsbD*, *PsbC*, *PsbB* as detected with *PsbA* being more expressed in more severe conditions, especially at Site 1 in March 2023, and at Site 2 in August 2023. At Site 3, the increased expression of PSII core genes *PsbD*, *PsbC*, *PsbB* was increased in August 2022 while the increase in *PsbA* showed weak trend toward October 2022 and August 2023. The expression of PSII core genes *PsbD*, *PsbC*, and *PsbB* was correlated positively with majority of small PSII subunits, with exception of *PsbX* which is more positive correlated to *PsbA*. The light harvesting complex genes *LhcA*, *LhcB* and *Lhcb4* had the tendency to increase in August 2022, and their expression was accompanied by increase in expression of the PSII small subunit *PsbK* as well as the expression of *Elip*.

However, different trends were observed between sites, probably due to the absence of March as the most severe conditions at Sites 2 and 3. The PSII small subunit *PsbS* expression increased toward October 2022 at Sites 2 and 3, contrary to Site 1 where this gene was more expressed in August 2022. The expression of the PSII small subunits *PsbX* and *PsbW* was also closely related at Sites 2 and 3, but this trend was not seen at Site 1. Finally, expression of all oxygen evolving complex genes was related tightly in Site 1 in October 2023, while at Sites 2 and 3, maximum expression of these genes occurred in August. The genes *PsbO* and *PsbU* were more expressed in August 2022, and genes *PsbQ* and *PsbP* in August 2023. he regulation of PSII assembly via Ycf48 was more prominent at Site 1 in October 2023, whereas it was more expressed at Sites 2 and 3 in August 2022. The gene *Cor413pm1* was expressed more at Site 1 in August 2023, while at Sites 2 and 3 in October 2022 and August 2023. At Sites 1 and 2, the *RbcS* expression was correlated to October 2022, contrasting with Site 3, where raised expression was related to August. Surprisingly, no transcripts of PSII assembly regulation gene *Ohp1* were detected at Site 3 (Supplement S11).

## 4 Discussion

### 4.1 Microclimate data

In general, the microclimate data obtained in this study did not differ from the average numbers for the Svalbard region (Láska *et al*. 2012; Norwegian Meteorological Institute 2023; UNIS weather stations 2023). According to the air temperature data, the coldest and warmest months (March and July, respectively) exhibited similar patterns and did not differ between the localities.

Regarding the soil temperature data, data of ground temperature from nearby Petunia Bay were similarly shown to be the lowest in “spring” months (April 2009 and March 2010) (Láska *et al*. 2012). However, according to the measured soil temperature data in this study, the coldest month was December 2022, probably due to a low level or absence of snow cover, which could act as thermal insulation uncoupling air temperature fluctuations from ground temperatures (Davey *et al*. 1992; Fahnestock *et al*. 1998; Morgner *et al*. 2010). Early and deep snow accumulation may even protect microbial populations from extreme and harmful freezing conditions, and potentially allows them to maintain some level of activity during winter in comparison to areas with delayed snow cover (Fahnestock *et al*. 1998; Morgner *et al*. 2010). Site 1, the lowest elevation site where snow probably accumulated later in winter, showed unexpectedly extreme conditions with the lowest soil temperatures and the high variability measured.

Site 2 appeared to be more humid than Site 3, which could be caused by the presence of clouds at this elevation (personal observation). However, this observation was not supported by the irradiance data.

### 4.2 Diversity of algae and cyanobacteria

Microbial phototrophs are essential for the process of photosynthesis, contributing a substantial portion of the Earth’s oxygen and aiding in the maintenance of atmospheric gas equilibrium. Photosynthetic microorganisms function as the main primary producers in the most extreme polar regions (Borowitzka *et al*. 2016; Pichrtová *et al*. 2020). This investigation corroborates preceding discoveries related to the community compositions of algae and cyanobacteria in analogous biocrusts present in polar or alpine environments, as detected through light microscopy (Čapková *et al*. 2016; Weber *et al*. 2016; Borchhardt *et al*. 2017b, 2017a) or examined using molecular data (Rippin *et al*. 2018a, 2018c; Pushkareva *et al*. 2024a). The diversity of microbial phototrophs in polar regions may be influenced by various environmental factors. For instance, the composition appears to be affected by the altitudinal gradient (Kotas *et al*. 2018), and different substrate types or snow cover dynamics further support distinct microbial communities (Zinger *et al*. 2009; Savaglia *et al*. 2024). However, the investigated sites did not differ in terms of community composition and the bedrock and soil composition, such as sandstones and siltstones, and soil types of silt loam and loam were relatively similar.

Detected eukaryotic algae belong according to molecular analyses mainly to the Trebouxiophyceae and Chlorophyceae groups, which appear to be typical inhabitants of biocrusts in Svalbard (Borchhardt *et al*. 2017a; Rippin *et al*. 2018a). Interestingly the same dominant groups are reported from Maritime Antarctica, where in contrast an abundance of Ulvophyceae (Pushkareva *et al*. 2024a) and Bacillariophyceae (Borchhardt *et al*. 2017b) are reported. However, several Icelandic biocrusts also showed even higher volumes of algae and a dominance of diatoms according to biovolume data (Pushkareva *et al*. 2021).

The cyanobacterial community data indicated a prevalence of filamentous groups belonging to the Nostocales, Pseudanabaenales, and Oscillatoriales, which is consistent with earlier findings from Arctic and Antarctic biocrusts (Rippin *et al*. 2018b; Pushkareva *et al*. 2024a). Filamentous cyanobacteria represent a characteristic group inhabiting polar terrestrial environments, showing high resistance to stressful conditions such as freezing or desiccation (Davey 1989; Hawes *et al*. 1992; Šabacká and Elster 2006; Tashyreva and Elster 2016).

### 4.3 Photosynthetic activity

In this study, the photosynthetic activity of biological soil crust in the High Arctic was evaluated using variable chlorophyll fluorescence measurement techniques, which are widely used for the evaluation of various stress and stress tolerance determination of plants and microalgae (Büchel and Wilhelm 1993; Maxwell and Johnson 2000; Consalvey *et al*. 2005; Yadav *et al*. 2023). Our observations confirmed diel periodical changes in photosynthetic activity, which is in agreement with previous findings measured on algae communities (Kvíderová *et al*. 2019), lichens (Sehnal *et al*. 2014), and even biocrusts in the Arctic (Sehnal *et al*. 2014; Pushkareva *et al*. 2017) or other arid and semi-arid temperate regions (Lan *et al*. 2012; Wu *et al*. 2013). However, a detailed comparison of the F_V_/F_M_ values could be complicated by different approaches of studies. First of all, F_V_/F_M_was measured on biocrusts after 15 minutes of dark acclimation, but in the other studies the parameter could also be measured after shorter acclimation (e.g. only 8 minutes) (Wu *et al*. 2013), longer (more than 20 minutes) (Lan *et al*. 2012) or even without any dark acclimation presented as the effective quantum yield Φ_PSII_ (Sehnal *et al*. 2014) depending on research tasks, experimental design, fluorescence measurement protocol used and evaluated organisms/consortia. However, from a comparison between 15 min dark acclimated F_V_/F_M_ and Φ_PSII_ values without dark acclimation, the results showed a strong positive correlation (Pushkareva *et al*. 2017).

More than duration of the dark adaptation period, the F_V_/F_M_ could be affected by stresses encountered *in situ*. Previous studies have shown that a decrease in photosynthetic activity can result from damage to the photosynthetic apparatus caused by freezing, irradiance, and/or desiccation (Davey, 1989; Hawes, 1990; Remias *et al*., 2018; Yoshida *et al*., 2020). The highest observed F_V_/F_M_ value in our measurements was 0.63, which is much higher than the highest value of 0.47 measured previously in a similar type of Svalbard biocrusts from Petunia Bay in Svalbard (Pushkareva *et al*. 2017), and slightly lower than the 0.7 reached in another similar study conducted idem (Sehnal *et al*. 2014) indicating relatively good physiological performance at all sites in summer. Surprisingly, similar values were detected even at Site 1 in early October 2022 when the environmental conditions were deteriorating, indicating thus good physiological state of photosynthetic microorganism at near-zero temperatures and in low light. Contrary in late October 2023, the stress conditions were more severe and led to significant decrease in F_V_/F_M_. Regardless the different F_V_/FM values in autumn in both years, the rETR_max_ was very reduced due to extreme low light conditions. Furthermore, a relation to water availability was previously suggested (Sehnal *et al*. 2014). However, in our study the influence of relative humidity could not be confirmed due to the small number of available data. Temperature has also been reported to play a role in cold environments. The increase in temperature caused a decrease in quantum yields in biocrusts (Sehnal *et al*. 2014; Pushkareva *et al*. 2017) and minor effects of temperature were reported from tidal flats dominated by *Vaucheria* sp. (Kvíderová *et al*. 2019). However, in this study, F_V_/F_M_ and temperature significantly correlated only at Site 1. Interestingly, the highest mean values of the F_V_/F_M_ were observed in the beginning of October, which is in agreement with the temperature and irradiance influence mentioned above. Detailed measurement focused on estimation if the compensation irradiance, i.e. when the photosynthesis is equal to respiration, should be performed to determine if the actual irradiance is sufficient for net primary production. Therefore, photophysiological data can provide valuable insight into the tolerance limits of algae and they could be used for better prediction of primary productivity in polar ecosystems.

Above abiotic conditions, the results could depend on dominant organisms. The measured maxima among different unstressed groups of algae vary: for example, diatoms reached 0.6 while the chlorophyte maximum was 0.8 (Büchel and Wilhelm 1993). Since the polar biological crusts are not homogeneous due to different nanoclimatic, edaphic and orographic conditions, there should be spots with different prevailing microorganisms. Considering the signal integration from whole during the fluorometer measurements, the F_V_/F_M_ could be lower due to inclusion of cyanobacteria-rich crusts or of non-photosynthetic areas like bare soil. Fluorescence imaging cameras should reveal this hidden physiological variability, and multispectral/hyperspectral images could be used for determination of the crust types (e.g. Rodríguez-Caballero *et al*. 2014). Furthermore, it has been suggested in the literature, that the increase in the photosynthetic activity in biocrusts could be related to the level of succession (Gypser *et al*. 2016; Pushkareva *et al*. 2017).

The selected OJIP parameters responded to seasonal changes. In mild conditions, the parameters related to photochemical quenching were creased. In more stressing conditions, the parameters reflecting non-photochemical energy dissipation rose, reflecting thus damage of the photosynthetic apparatus and increased need to get rid of the excess of light energy (Roháček *et al*. 2008). The decreased number of active reaction centres led to increased energy fluxes in less favourable conditions. It has previously been shown that the diurnal courses of some OJIP parameters (mostly fluxes per active reaction centres), significantly correlated with temperature and varied between different types of biocrusts (Pushkareva *et al*. 2017). This agrees with the results of ordination analyses in this study, where site appeared to be the most influential.

For both seasons of the year, photosynthetic activity mostly followed the irradiance changes, to which F_V_/F_M_ negatively and rETR_max_ positively correlated. In the summer cycles, the maximum FV/F_M_ was observed around midnight and the „night“ hours when the lower irradiances encountered, while the maximum rETR_max_ was detected around noon and „day“ hours, potentially indicating only low photoinhibition. Such midday depression of F_V_/F_M_ or Φ_PSII_ was also previously detected in polar (Sehnal *et al*. 2014; Pushkareva *et al*. 2017) and temperate desert biocrusts (Lan *et al*. 2012; Wu *et al*. 2013), but this depression not always leads to a reduction in photosynthetic performance in mid-day or early afternoon which was observed by biocrusts (Pushkareva *et al*. 2017).

### 4.4 Recovery of photosynthetic activity after winter biocrusts thawing

The rapid increase in quantum yield observed in thawed biocrusts during the winter season indicates that they possess the capacity to restore photosynthetic activity at a rapid rate following the attainment of an optimal environmental condition. Similarly, it has previously been reported that polar cryptogams (mosses and lichens) can recover their photosynthetic activity in a matter of minutes or hours (Schlensog *et al*. 2004). The rapid recovery of photosynthetic activity probably crucial for survival in unstable polar environments.

Surprisingly, the results of photosynthetic activity measured in late October (23/10/2023) indicated biocrust activity even at subzero temperatures. The values were naturally low due to the stressful conditions typical of the Arctic autumn environment. However, differences in the daily cycle remained evident. Cyanobacteria and algae have already been shown to perform photosynthetic activity at temperatures down to −7 °C (Davey 1989) and general soil respiration activity has been evidenced at −12 °C (Elberling 2007). Polar cryptogams, such as lichens, have been demonstrated to retain their activity at subzero temperatures (Kappen *et al*. 1996; Hájek *et al*. 2016, 2021; Barták *et al*. 2021) even at −17 °C (Kappen *et al*. 1996). However, the spring and autumn months are likely to be the most significant periods for their primary production (Schroeter *et al*. 1995), given that their continued existence is contingent upon water availability (Kappen *et al*. 1990; Hovenden *et al*. 1994; Laguna-Defior *et al*. 2016). Vascular plants at high latitudes exhibit high freezing tolerance; however, in contrast, they are not actively photosynthesizing during the winter, resulting in a relatively short vegetative season (Kappen 1993; Laurila *et al*. 2001; Arndal *et al*. 2009). Although polar biocrusts exhibit lower photosynthetic rates compared to vascular plants, they may benefit from a more consistent water supply from soil, in contrast to lichens. This together with their ability to quickly restore photosynthetic activity allows them to serve as crucial contributors to carbon fixation during periods of inactivity for other organisms, thus underscoring their role within polar ecosystems.

### 4.5 Photosynthesis-related genes

Data for photosynthesis-related transcripts indicate that photosynthesis-related genes are expressed in biocrusts throughout the year, even in March. It suggests that biocrusts are always ready to engage in photosynthetic activity even during the winter months, although at reduced levels compared to other seasons. The results are also consistent with the photosynthetic activity observed in the biocrusts during the autumn monitoring period and immediately after thawing in the winter experiment. In winter, photosynthesis was probably driven by the availability of ambient light during the sampling process, which appeared sufficient to stimulate metabolic processes despite extreme cold. In fact, the soil samples taken during the sampling period were frozen. Therefore, the observed transcriptome probably reflects the levels present when the biocrust samples were last frozen before March. It should be noted that the microbial communities within these biocrusts have been shown to be permanently well prepared to cope with the extreme environment of the Arctic (Pushkareva *et al*. 2024b). To investigate this further, we employed the cold shock marker *Cor413pm1* (Hu *et al*. 2021), which we could detect in all samples. No significant changes in the expression level among sites nor seasons were observed in biocrusts, supporting the earlier results of Pushkareva *et al*. (2024b). However, a closer look at the metatranscriptomic data revealed that the biocrusts still needed to acclimate to the different seasons and locations. The expression levels of *PsbA* and *RbcS* changed dramatically, especially in the March samples, suggesting strong photoinhibition by photodamage that led to increased *PsbA* turnover (Andersson and Aro 2001; Mulo *et al*. 2012), which is reflected in our transcriptomic data. In *Chlamydomonas* acclimation to high light suppressed *RbcL* expression temporarily (Shapira *et al*. 1997), suggesting that the observed decrease of *RbcS* might be due to acclimation to light. The need to acclimate to light during winter is also supported by the seasonal regulation of Ohp1 and ELIP proteins, which have been shown to be important during light acclimation (Adamska 1997; Jansson *et al*. 2000). Why other proteins of the PSII also show differential gene expression related to site and season, is currently not clear. Clearly, more studies on this interesting phenomenon are needed to solve these questions.

### 4.6 Photosynthetic activity and gene expression

The variable chlorophyll fluorescence originates in PSII predominantly (Krause and Weis 1991). Therefore, any changes in PSII function or structure could be reflected in the fluorescence signal, and *vice versa*, the change in fluorescence parameters could precede any observable damage and/or stress response (Solovchenko *et al*. 2022). The summer season should be less stressful than winter, therefore increased photochemical quenching reflected especially by increase in FV/F_M_ and φ_ET2o_ should be expected. Since summer is also the main growth season in the Polar Regions, there is high demand for synthesis of new PSIIs as well. Indeed, higher F_V_/F_M_ and φ_ET2o_ were observed in summer, especially in August 2022, accompanied by high levels of transcripts for PSII core proteins *PsbD*, *PsbC* and *PsbB*, majority of minor subunits, oxygen evolving complex proteins and light harvesting complex proteins. In more stressing conditions, in autumn and winter, and to some extent even in August 2023, the increased non-photochemical quenching expressed as φ_Do_, and hence decline in photochemical quenching, indicating stress conditions was observed. The increased *PsbA* expression indicating rapid protein turnover due to its damage was detected as well (Giardi *et al*. 1997). Increased expression of by *PsbS*, small PSII subunit participating in non-photochemical quenching (Morosinotto and Bassi 2014), and *PsbX*, involved in the binding and/or turnover of quinones at the Q_B_ site of PSII to maintain efficient electron transport also confirmed stress conditions (Biswas and Eaton-Rye 2022). Although the electron transport remains functional even in sub-zero temperatures (Davey 1989), the low temperatures lead to delay in the rates of electron transfer in PSII resulting in photooxidative damage (Pospíšil 2016). The high amount of *PsbA* transcripts was indeed detected (see discussion above), and in case of fluorescence parameters, low F_V_/F_M_ and φET2o together with increased φ_Do_ should be detected. The V_J_ should be increased due to delay in electron transport to Q_B_ and the time to reach the maximum fluorescence should be longer. More detailed fluorescence protocols including quenching analysis and rapid light curves should be implemented together with transcriptome analysis to reveal interplay between energetic metabolism and gene expression, especially in extreme conditions.

## 5 Conclusion

The findings of this study indicate that diurnal variation in the photosynthetic activity of biological soil crusts at three sites, which exhibited a relatively comparable microbial phototrophic composition, is mainly influenced by irradiance during the summer and autumn seasons of 2022–2023. No significant differences were observed in the diurnal variation of photosynthetic activity between the sites. Of particular interest is the markedly elevated level of activity observed in the initial days of October 2022, which persisted at temperatures below zero in the latter days of October 2023. Furthermore, winter-thawed biocrusts exhibited the capacity to restore photosynthesis rapidly following thawing. These findings are also supported by the metagenomic and metatranscriptomic data of microbial phototrophs and photosynthesis-related genes and provide valuable information on the behaviour of the studied organisms and emphasise the importance of environmental factors such as temperature and irradiance in influencing their activity levels.

## Data availability statement

The data that support the findings of this study are available on request from the corresponding author. Raw reads were submitted to the Sequence Read Archive under the project PRJNA1124630.

## Author contributions

Conceptualization: BB, JE. Methodology: EH, BB, JE, JK, EP. Formal analysis: EH, EP. Investigation: EH, EP. Visualisation: EH, JK. Writing – Original Draft: EH. Writing – Review & Editing: EH, BB, JE, JK, EP. Funding acquisition: BB, JE.

## Funding

The project was supported by the Grant Agency of the Czech Republic (22-08680L), the Institute of Botany of the Czech Academy of Sciences (RVO67985939), the Charles University Research Centre program (UNCE/24/SCI/006), and by the German Research Foundation (BE1779/25).

## Acknowledgments

The authors would like to thank the Czech Arctic Research Station of the University of South Bohemia in České Budějovice for providing their facilities in Svalbard. Alžběta Paterová, Elisa Frank, Anastasiia Kolomiiets, Oleksandr Bren and Paul Geroldinger are acknowledged for field assistance, and Viktorie Brožová for help with determination of vegetation. Jana Kvíderová is grateful for support from the University of West Bohemia. Tyler J. Kohler is thanked for proofreading the manuscript.

## Supplementary material

Supplement S1. Geographic, vegetation, and geological characteristics of the investigated sites near Longyearbyen, West Spitsbergen, Svalbard.

Supplement S2. Photographs showing a) the installation of Petri dish (left) and plastic bowl (right), b) the dark acclimation process.

Supplement S3. The list of used OJIP parameters and their physiological meanings adopted from Stirbet *et al*. 1998; Strasser *et al*. 2004. RC (reaction centre), PSII (photosystem II), Q_A_ (primary acceptor plastoquinone), Q_B_ (secondary acceptor plastoquinone).

Supplement S4. Overview of microclimate data including month means, minimum and maximum values.

Supplement S5. The diurnal change of values of environmental parameters, maximum quantum yield (F_V_/F_M_; mean ± s.d.) and maximum possible relative electron transport rate (rETR_max_; mean ± s.d.) during in situ measurement of photosynthetic activity. The presence of diurnal changes was tested by one-way ANOVA. Abbreviations: ANOVA – one-way ANOVA, n – number of cases, n.m. – not measured, PAR – photosynthetically active radiation, RH – relative air humidity, T_air_ – air temperature, T_soil_ – soil temperature. The statistically significant differences are marked in bold. Data used: averages per Petri dish and bowl.

Supplement S6. The diurnal changes of the photosynthetic (F_V_/F_M_ and rETR_max_; mean ± s.d., for n refer to Supplement S5) and environmental parameters (air and soil temperature, T_air_, T_soil_; photosynthetically active radiation, PAR; relative humidity, RH) at all the experimental sites in the studied periods in August 2022 and 2023.

Supplement S7. Correlations of the F_V_/F_M_ and rETR_max_ measured during diurnal cycles study with environmental data for the summer and autumn seasons 2022 and 2023. The statistically significant correlations are marked in bold.

Supplement S8. The changes of effective quantum yield (Ф_PSII_; mean ± s.d., n = 12) during recovery of photosynthetic activity in winter. The statistically significant differences were tested using Repeated Measures Analysis of Variance (RM ANOVA; n *=* 12). The letter in upper case indicates homologous groups recognized by Tukey HSD test for P = 0.05.

Supplement S9. Relative abundances of photosynthesis-related transcripts (expressed in percentage of FPKM, fragments per kilobase of transcript per million fragments sequenced, mean ± s.d.) per study site and sampling season.

Supplement S10. Results of two-factor ANOVA (n_(Aug22)_ = 5, n_(Oct22)_ = 2, n_(Mar23, Aug23)_ = 4) assessing the impact of site (Site 1 × Site 2 × Site 3) and sampling season (Aug22 × Oct22 × Mar23 × Aug23) on photosynthesis- and stress-related transcripts represented by FPKM numbers (fragments per kilobase of transcript per million fragments sequenced).

Supplement S11. RDA analyses showing correlation among relative abundances of photosynthesis-related transcripts (explained variables: relative abundances of photosynthesis-related genes; arrows) and environmental parameters (explaining variables: sampling season; red symbols) and separation of gene expression at individual sites. The total variation is 480 (Site 1) / 320 (Site 2) / 341 (Site 3), explanatory variables account for 51.28 % / 37.82 % / 41.73 % of explained variation. Monte Carlo Permutation test results: P = 0.002 / P = 0.004/ P = 0.008, pseudo-F = 1.9 / 1.7 / 1.7 (first axis); P = 0.002 / P = 0.022 / P = 0.002, pseudo-F = 1.9 / 1.7 / 1.7 (all axes).

## Notes

### Competing Interest Statement

The authors have declared no competing interest.

