## Supplements S1-S11 for "Summer and autumn photosynthetic activity in High Arctic biological soil crusts and their winter recovery"

### Supplementary material

Hejduková E., Pushkareva E., Kvíderová J., Becker B., Elster J.

**Supplement S5**. The diurnal change of values of environmental parameters, maximum quantum yield ($F_V/F_M$; mean ± s.d.) and maximum possible relative electron transport rate ($rETR_{max}$; mean ± s.d.) during *in situ* measurement of photosynthetic activity. The presence of diurnal changes was tested by one-way ANOVA. Abbreviations: ANOVA – one-way ANOVA, n – number of cases, n.m. – not measured, PAR – photosynthetically active radiation, RH – relative air humidity, $T_{air}$ – air temperature, $T_{soil}$ – soil temperature. The statistically significant differences are marked in bold. Data used: averages per Petri dish and bowl. ........ 9

**Supplement S11**. RDA analyses showing correlation among relative abundances of photosynthesis-related transcripts (explained variables: relative abundances of photosynthesis-related genes; arrows) and environmental parameters (explaining variables: sampling season; red symbols) and separation of gene expression at individual sites. The total variation is 480 (Site 1) / 320 (Site 2) / 341 (Site 3), explanatory variables account for 51.28 % / 37.82 % / 41.73 % of explained variation. Monte Carlo Permutation test results: P = 0.002 / P = 0.004/ P = 0.008, pseudo-F = 1.9 / 1.7 / 1.7 (first axis); P = 0.002 / P = 0.022 / P = 0.002, pseudo-F = 1.9 / 1.7 / 1.7 (all axes). ........ 16

**Supplement S1**. Geographic, vegetation, and geological characteristics of the investigated sites near Longyearbyen, West Spitsbergen, Svalbard.

| Site | GPS coordinates | Elevation | Site length | Site width | Site area | Vegetation | Bedrock | Snow depth (March 2023) |
|---|---|---|---|---|---|---|---|---|
| **Site 1** | 78°13'11.5"N 15°19'54.9"E | 47 m a.s.l. | 9.35 m | 5.75 m | 53.76 $m^2$ | *Saxifraga cespitosa*, *Oxyria digyna*, *Minuartia biflora*, *Salix polaris*, *Silene acaulis*, *Luzula* sp., mosses | sandstone, siltstone, shale intercalations, locally coal seams (close to the Cretaceous-Tertiary boundary) | 40–78 cm |
| **Site 2** | 78°09'20.3"N 16°01'52.7"E | 409 m a.s.l. | 8.41 m | 2.51 m | 21.11 $m^2$ | *Polytrichum* sp., *Luzula confusa*, *lichens*, *Cerastium arcticum*, *Saxifraga cespitosa*, liverwort | sandstone, siltstone, shale intercalations, locally coal seams (Cretaceous) | 58–68 cm |
| **Site 3** | 78°08'47.4"N 16°02'21.5"E | 519 m a.s.l. | 8.30 m | 7.65 m | 63.50 $m^2$ | *Aulacomnium turgidum*, *Saxifraga cernua*, *Phippsia algida*, lichens | sandstone, siltstone and shale | 116–156 cm |

**Supplement S2**. Photographs showing a) the installation of Petri dish (left) and plastic bowl (right), b) the dark acclimation process.

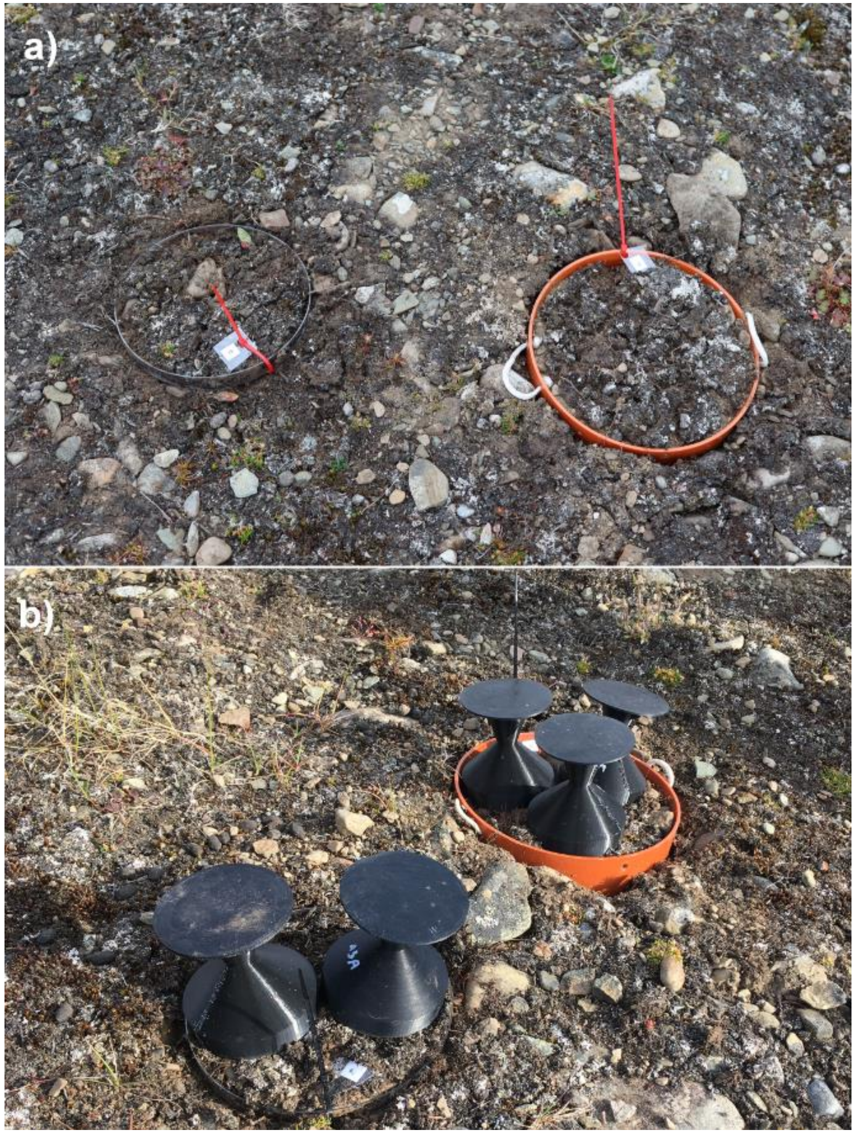

**Supplement S3**. The list of used OJIP parameters and their physiological meanings adopted from Stirbet et al. 1998; Strasser et al. 2004. RC (reaction centre), PSII (photosystem II), $Q_A$ (primary acceptor plastoquinone), $Q_B$ (secondary acceptor plastoquinone).

| Parameter | FluorPen (export) | Physiological meaning | In stressful conditions | Theoretical range |
|---|---|---|---|---|
| **Technical parameters** | | | | |
| $M_0$ | Mo | Maximum rate of accumulation of closed RCs at the beginning of fluorescence rise | Increase | 0–4 (theoretical for $F_{300}=F_M=6.25\times F_0$ and maximum $F_V/F_M=0.84$) |
| $V_J$ | Vj | Normalized fluorescence intensity at 2 ms (J-step) | Increase | 0–1 |
| $V_I$ | Vi | Normalized fluorescence intensity at 30 ms (I-step)* | Increase | 0–1 |
| **Quantum yields** | | | | |
| $F_V/F_M=\varphi_{Po}$ | Phi_Po | Maximum quantum yield of primary photochemistry of PSII | Decrease | 0–0.84 |
| $\varphi_{ET2o}$ | Phi_Eo | Quantum yield of electron transport flux from $Q_A$ to $Q_B$ | Decrease | 0–0.84 |
| $\varphi_{Do}$ | Phi_Do | Quantum yield of energy dissipation | Increase | 0.16–1 (theoretical for maximum $F_V/F_M=0.84$) |
| **Efficiencies/Probabilities** | | | | |
| $\psi_{ET2o}$ | Psi_0 | Efficiency/probability that electron trapped by PSII will be transferred to $Q_A$ | Decrease | 0–1 |
| **Fluxes through active RC** | | | | |
| $J_0^{ABS}/RC$ | ABS/RC | Average absorbed photon flux per active RC | Increase/decrease** | 0–4.85 |
| $J_0^{TR}/RC$ | TRo/RC | Maximum trapped electron flux per active RC | Increase/decrease** | 0–4 |
| $J_0^{ET2}/RC$ | ETo/RC | Electron transport flux from $Q_A$ to $Q_B$ per active RC | Increase/decrease** | 0–4 |
| $J_0^{DI}/RC$ | DIo/RC | Energy flux dissipated as heat | Increase/decrease** | 0–∞ |

* depending on the inflection point position, 60 ms timing could be used
** depending on active RC number

**Supplement S4.** Overview of microclimate data including month means, minimum and maximum values.

| | Site | Month, Year | n | Month mean | | Day mean minimum | | Day mean maximum | | Minimum measured | | Maximum measured | |
|---|---|---|---|---|---|---|---|---|---|---|---|---|---|
| | | | | Mean | s.d. | Min | Day | Max | Day | Min | Day, Hour | Max | Day, Hour |
| **Air temperature (°C)** | | Aug 2022 | 636 | 6.26 | 2.69 | 3.55 | 8.30.22 | 10.00 | 8.6.22 | 0.66 | 8.28.22 6:00 | 14.00 | 8.11.22 10:00 |
| | | Sep 2022 | 720 | 2.45 | 3.14 | −3.67 | 9.19.22 | 7.23 | 9.3.22 | −5.62 | 9.19.22 6:00 | 12.13 | 9.26.22 12:00 |
| | | Oct 2022 | 744 | −2.37 | 4.25 | −9.94 | 10.26.22 | 3.52 | 10.8.22 | −11.18 | 10.26.22 10:00 | 5.06 | 10.7.22 7:00 |
| | | Nov 2022 | 720 | −1.72 | 3.51 | −8.37 | 11.26.22 | 4.16 | 11.29.22 | −9.87 | 11.26.22 23:00 | 5.00 | 11.29.22 1:00 |
| | | Dec 2022 | 744 | −8.76 | 6.55 | −18.86 | 12.14.22 | 4.21 | 12.2.22 | −20.30 | 12.14.22 3:00 | 5.30 | 12.2.22 14:00 |
| | | Jan 2023 | 744 | −4.39 | 3.78 | −12.73 | 1.27.23 | 0.15 | 1.1.23 | −15.42 | 1.27.23 5:00 | 3.42 | 1.23.23 5:00 |
| | | Feb 2023 | 672 | −6.94 | 4.41 | −13.90 | 2.20.23 | −0.11 | 2.24.23 | −15.30 | 2.19.23 8:00 | 1.92 | 2.14.23 20:00 |
| | Site 1 | Mar 2023 | 744 | −13.66 | 4.26 | −19.46 | 3.16.23 | −3.92 | 3.2.23 | −22.33 | 3.21.23 6:00 | −2.80 | 3.2.23 13:00 |
| | | Apr 2023 | 720 | −5.43 | 5.07 | −15.16 | 4.4.23 | 1.20 | 4.15.23 | −17.18 | 4.4.23 20:00 | 4.59 | 4.15.23 4:00 |
| | | May 2023 | 744 | −2.05 | 3.64 | −12.21 | 5.2.23 | 2.93 | 5.23.23 | −14.97 | 5.2.23 3:00 | 6.26 | 5.23.23 11:00 |
| | | Jun 2023 | 720 | 3.24 | 3.26 | −0.02 | 6.2.23 | 9.65 | 6.21.23 | −3.63 | 6.2.23 6:00 | 12.89 | 6.21.23 12:00 |
| | | Jul 2023 | 744 | 9.67 | 2.34 | 6.76 | 7.1.23 | 13.27 | 7.14.23 | 4.97 | 7.6.23 6:00 | 16.25 | 7.5.23 10:00 |
| | | Aug 2023 | 744 | 8.03 | 1.95 | 5.02 | 8.23.23 | 11.82 | 8.4.23 | 4.17 | 8.23.23 5:00 | 15.92 | 8.3.23 14:00 |
| | | Sep 2023 | 720 | −0.19 | 4.43 | −8.69 | 9.27.23 | 6.13 | 9.5.23 | −9.41 | 9.27.23 6:00 | 8.29 | 9.8.23 15:00 |
| | | Oct 2023 | 543 | −4.40 | 2.69 | −9.33 | 10.22.23 | 0.32 | 10.19.23 | −10.15 | 10.22.23 2:00 | 2.34 | 10.19.23 4:00 |
| | | Aug 2022 | 612 | 3.66 | 2.65 | 0.08 | 8.27.22 | 7.41 | 8.6.22 | −2.63 | 8.28.22 0:00 | 11.91 | 8.12.22 13:00 |
| | | Sep 2022 | 720 | 0.89 | 2.96 | −3.56 | 9.19.22 | 5.43 | 9.2.22 | −7.07 | 9.19.22 5:00 | 9.21 | 9.26.22 12:00 |
| | | Oct 2022 | 744 | −5.21 | 4.18 | −12.04 | 10.26.22 | 0.95 | 10.1.22 | −12.71 | 10.25.22 17:00 | 2.29 | 10.1.22 15:00 |
| | | Nov 2022 | 720 | −3.71 | 3.88 | −10.78 | 11.26.22 | 1.84 | 11.29.22 | −11.97 | 11.26.22 18:00 | 2.65 | 11.28.22 21:00 |
| | | Dec 2022 | 744 | −11.83 | 6.89 | −22.10 | 12.14.22 | 1.43 | 12.2.22 | −23.72 | 12.13.22 15:00 | 2.51 | 12.2.22 13:00 |
| | | Jan 2023 | 744 | −7.45 | 4.01 | −15.79 | 1.27.23 | −2.30 | 1.5.23 | −18.80 | 1.26.23 22:00 | 1.23 | 1.23.23 5:00 |
| | | Feb 2023 | 672 | −9.91 | 4.62 | −17.91 | 2.2.23 | −3.46 | 2.24.23 | −19.86 | 2.2.23 20:00 | −0.13 | 2.14.23 21:00 |
| | Site 2 | Mar 2023 | 744 | −16.63 | 4.76 | −23.49 | 3.21.23 | −5.58 | 3.2.23 | −26.91 | 3.21.23 2:00 | −4.51 | 3.2.23 0:00 |
| | | Apr 2023 | 720 | −7.23 | 6.08 | −18.78 | 4.4.23 | 2.55 | 4.15.23 | −19.92 | 4.4.23 17:00 | 4.80 | 4.14.23 21:00 |
| | | May 2023 | 744 | −4.37 | 4.35 | −16.51 | 5.1.23 | 1.94 | 5.23.23 | −18.41 | 5.1.23 22:00 | 4.26 | 5.23.23 10:00 |
| | | Jun 2023 | 720 | 2.25 | 3.11 | −2.16 | 6.2.23 | 8.85 | 6.21.23 | −3.41 | 6.2.23 2:00 | 10.31 | 6.21.23 11:00 |
| | | Jul 2023 | 744 | 7.03 | 2.50 | 4.14 | 7.17.23 | 12.55 | 7.6.23 | 2.41 | 7.17.23 5:00 | 13.69 | 7.6.23 15:00 |
| | | Aug 2023 | 744 | 5.73 | 2.46 | 2.40 | 8.21.23 | 10.75 | 8.5.23 | 1.43 | 8.21.23 6:00 | 14.98 | 8.4.23 14:00 |
| | | Sep 2023 | 720 | −2.15 | 3.90 | −8.98 | 9.27.23 | 4.99 | 9.4.23 | −9.91 | 9.27.23 3:00 | 7.17 | 9.4.23 12:00 |
| | | Oct 2023 | 563 | −6.88 | 2.44 | −11.57 | 10.22.23 | −1.73 | 10.19.23 | −12.80 | 10.22.23 15:00 | 0.49 | 10.19.23 13:00 |
| | | Aug 2022 | 576 | 1.27 | 2.08 | −2.08 | 8.27.22 | 4.81 | 8.10.22 | −3.70 | 8.28.22 1:00 | 6.00 | 8.10.22 9:00 |
| | | Sep 2022 | 720 | −0.52 | 2.98 | −5.23 | 9.17.22 | 4.28 | 9.3.22 | −6.91 | 9.13.22 1:00 | 5.54 | 9.3.22 5:00 |
| | | Oct 2022 | 744 | −6.13 | 3.88 | −11.74 | 10.25.22 | 0.12 | 10.1.22 | −12.39 | 10.19.22 3:00 | 1.74 | 10.1.22 1:00 |
| | | Nov 2022 | 720 | −4.31 | 3.48 | −9.99 | 11.9.22 | 0.91 | 11.29.22 | −11.42 | 11.17.22 5:00 | 1.97 | 11.2.22 0:00 |
| | Site 3 | Dec 2022 | 744 | −12.54 | 6.71 | −22.37 | 12.13.22 | 0.47 | 12.2.22 | −23.90 | 12.14.22 8:00 | 1.32 | 12.2.22 12:00 |
| | | Jan 2023 | 744 | −8.36 | 3.92 | −16.76 | 1.26.23 | −3.47 | 1.5.23 | −18.09 | 1.26.23 16:00 | 0.66 | 1.23.23 3:00 |
| | | Feb 2023 | 672 | −10.50 | 4.40 | −17.82 | 2.19.23 | −4.58 | 2.24.23 | −19.37 | 2.18.23 21:00 | −0.96 | 2.14.23 19:00 |
| | | Mar 2023 | 744 | −17.69 | 4.65 | −24.37 | 3.21.23 | −6.90 | 3.2.23 | −26.54 | 3.21.23 4:00 | −4.78 | 3.2.23 0:00 |
| | | Apr 2023 | 720 | −8.18 | 6.00 | −20.10 | 4.21.23 | 1.45 | 4.15.23 | −20.95 | 4.21.23 22:00 | 3.74 | 4.14.23 19:00 |

| | | | | | | | | | | | | |
|---|---|---|---|---|---|---|---|---|---|---|---|---|
| | | May 2023 | 744 | −5.72 | 4.18 | −17.98 | 5.1.23 | 0.09 | 5.23.23 | −18.53 | 5.1.23 4:00 | 2.22 | 5.20.23 14:00 |
| | | Jun 2023 | 720 | 0.82 | 3.56 | −4.29 | 6.1.23 | 9.04 | 6.21.23 | −5.26 | 6.2.23 1:00 | 10.46 | 6.21.23 8:00 |
| | | Jul 2023 | 744 | 6.27 | 2.53 | 3.10 | 7.3.23 | 10.28 | 7.6.23 | 1.46 | 7.3.23 0:00 | 11.70 | 7.6.23 14:00 |
| | | Aug 2023 | 744 | 5.45 | 2.72 | 1.85 | 8.21.23 | 11.12 | 8.4.23 | 0.68 | 8.21.23 4:00 | 14.06 | 8.4.23 14:00 |
| | | Sep 2023 | 720 | −2.73 | 3.88 | −8.89 | 9.27.23 | 4.75 | 9.4.23 | −9.63 | 9.27.23 4:00 | 6.34 | 9.4.23 12:00 |
| | | Oct 2023 | 564 | −7.74 | 2.55 | −12.59 | 10.22.23 | −2.53 | 10.19.23 | −13.86 | 10.21.23 20:00 | 0.04 | 10.19.23 13:00 |

| | Site | Month | n | Month mean | | Day mean minimum | | Day mean maximum | | Minimum measured | | Maximum measured | |
|---|---|---|---|---|---|---|---|---|---|---|---|---|---|
| | | | | Mean | s.d. | Min | Day | Max | Day | Min | Day, Hour | Max | Day, Hour |
| **Soil temperature (°C)** | Site 1 | Aug 2022 | 636 | 6.48 | 3.01 | 2.73 | 8.28.22 | 10.59 | 8.5.22 | 0.16 | 8.28.22 8:00 | 15.01 | 8.12.22 17:00 |
| | | Sep 2022 | 720 | 2.32 | 2.47 | −0.66 | 9.19.22 | 6.47 | 9.3.22 | −0.86 | 9.19.22 20:00 | 7.84 | 9.2.22 15:00 |
| | | Oct 2022 | 744 | −2.97 | 3.79 | −8.67 | 10.26.22 | 2.43 | 10.1.22 | −9.50 | 10.27.22 2:00 | 3.44 | 10.1.22 12:00 |
| | | Nov 2022 | 720 | −2.19 | 2.30 | −6.57 | 11.10.22 | 0.76 | 11.21.22 | −8.17 | 11.12.22 4:00 | 1.23 | 11.21.22 4:00 |
| | | Dec 2022 | 744 | −7.97 | 5.11 | −16.26 | 12.25.22 | 0.68 | 12.2.22 | −17.37 | 12.25.22 17:00 | 1.19 | 12.3.22 9:00 |
| | | Jan 2023 | 744 | −4.21 | 0.75 | −5.87 | 1.27.23 | −3.06 | 1.23.23 | −6.16 | 1.27.23 15:00 | −2.57 | 1.23.23 8:00 |
| | | Feb 2023 | 672 | −4.68 | 1.00 | −7.37 | 2.4.23 | −3.81 | 2.27.23 | −7.55 | 2.4.23 15:00 | −3.40 | 2.6.23 18:00 |
| | | Mar 2023 | 744 | −6.32 | 1.13 | −7.88 | 3.27.23 | −4.20 | 3.3.23 | −7.90 | 3.27.23 12:00 | −4.16 | 3.3.23 17:00 |
| | | Apr 2023 | 720 | −4.32 | 1.12 | −6.71 | 4.1.23 | −3.15 | 4.20.23 | −6.85 | 4.1.23 0:00 | −2.41 | 4.9.23 21:00 |
| | | May 2023 | 744 | −2.27 | 1.47 | −3.74 | 5.5.23 | −0.19 | 5.30.23 | −3.75 | 5.5.23 17:00 | −0.18 | 5.31.23 19:00 |
| | | Jun 2023 | 720 | 3.82 | 4.53 | −0.20 | 6.1.23 | 12.30 | 6.21.23 | −0.20 | 6.1.23 11:00 | 18.26 | 6.21.23 16:00 |
| | | Jul 2023 | 744 | 11.90 | 3.10 | 9.02 | 7.11.23 | 17.19 | 7.5.23 | 6.65 | 7.3.23 5:00 | 24.45 | 7.5.23 17:00 |
| | | Aug 2023 | 744 | 8.27 | 1.95 | 6.18 | 8.21.23 | 11.80 | 8.3.23 | 4.48 | 8.26.23 6:00 | 14.64 | 8.3.23 14:00 |
| | | Sep 2023 | 720 | 0.40 | 3.44 | −6.19 | 9.27.23 | 5.71 | 9.5.23 | −6.66 | 9.28.23 0:00 | 6.62 | 9.5.23 15:00 |
| | | Oct 2023 | 543 | −4.53 | 2.21 | −9.31 | 10.23.23 | −0.38 | 10.19.23 | −9.90 | 10.23.23 8:00 | 0.08 | 10.19.23 13:00 |
| | Site 2 | Aug 2022 | 612 | 4.05 | 2.79 | 0.53 | 8.28.22 | 8.70 | 8.7.22 | −1.21 | 8.28.22 3:00 | 11.13 | 8.7.22 11:00 |
| | | Sep 2022 | 720 | 0.94 | 1.89 | −1.71 | 9.22.22 | 4.96 | 9.2.22 | −2.02 | 9.22.22 8:00 | 6.50 | 9.2.22 16:00 |
| | | Oct 2022 | 744 | −3.89 | 3.19 | −9.24 | 10.26.22 | 0.10 | 10.3.22 | −9.79 | 10.21.22 12:00 | 0.63 | 10.3.22 13:00 |
| | | Nov 2022 | 720 | −2.97 | 1.83 | −8.07 | 11.26.22 | −0.53 | 11.19.22 | −9.16 | 11.26.22 20:00 | −0.33 | 11.20.22 5:00 |
| | | Dec 2022 | 744 | −9.59 | 4.12 | −15.48 | 12.25.22 | −0.80 | 12.3.22 | −15.83 | 12.25.22 14:00 | −0.46 | 12.3.22 16:00 |
| | | Jan 2023 | 744 | −6.29 | 0.38 | −6.97 | 1.19.23 | −5.64 | 1.15.23 | −7.16 | 1.1.23 0:00 | −5.63 | 1.15.23 19:00 |
| | | Feb 2023 | 672 | −6.45 | 0.57 | −7.93 | 2.5.23 | −5.83 | 2.27.23 | −7.98 | 2.5.23 2:00 | −5.82 | 2.27.23 12:00 |
| | | Mar 2023 | 744 | −8.14 | 1.12 | −9.73 | 3.27.23 | −6.20 | 3.1.23 | −9.78 | 3.27.23 19:00 | −6.06 | 3.1.23 0:00 |
| | | Apr 2023 | 720 | −7.20 | 0.95 | −8.95 | 4.1.23 | −5.77 | 4.18.23 | −9.10 | 4.1.23 0:00 | −5.75 | 4.18.23 17:00 |
| | | May 2023 | 744 | −4.33 | 2.10 | −6.85 | 5.4.23 | −0.90 | 5.24.23 | −6.86 | 5.4.23 6:00 | −0.87 | 5.24.23 12:00 |
| | | Jun 2023 | 720 | 0.76 | 3.25 | −1.94 | 6.3.23 | 10.74 | 6.30.23 | −1.97 | 6.4.23 13:00 | 15.34 | 6.30.23 15:00 |
| | | Jul 2023 | 744 | 8.84 | 2.76 | 5.41 | 7.11.23 | 14.23 | 7.7.23 | 3.54 | 7.4.23 4:00 | 17.57 | 7.7.23 13:00 |
| | | Aug 2023 | 744 | 6.73 | 2.10 | 4.45 | 8.29.23 | 11.08 | 8.5.23 | 3.05 | 8.17.23 1:00 | 12.22 | 8.4.23 17:00 |
| | | Sep 2023 | 720 | −1.34 | 3.62 | −8.56 | 9.27.23 | 4.35 | 9.4.23 | −9.05 | 9.28.23 0:00 | 5.34 | 9.4.23 12:00 |
| | | Oct 2023 | 563 | −5.19 | 1.55 | −8.82 | 10.10.23 | −3.08 | 10.20.23 | −9.30 | 10.10.23 23:00 | −3.02 | 10.20.23 5:00 |
| | Site 3 | Aug 2022 | 576 | 2.95 | 2.39 | −0.15 | 8.28.22 | 7.64 | 8.8.22 | −1.40 | 8.28.22 5:00 | 9.97 | 8.9.22 16:00 |

| | | | | | | | | | | | | | |
|---|---|---|---|---|---|---|---|---|---|---|---|---|---|
| | | Sep 2022 | 720 | 0.80 | 1.59 | −1.51 | 9.22.22 | 4.43 | 9.3.22 | −1.94 | 9.22.22 7:00 | 5.94 | 9.2.22 14:00 |
| | | Oct 2022 | 744 | −1.38 | 0.98 | −2.99 | 10.21.22 | 0.08 | 10.1.22 | −3.08 | 10.21.22 14:00 | 0.61 | 10.1.22 15:00 |
| | | Nov 2022 | 720 | −0.97 | 0.47 | −1.83 | 11.27.22 | −0.24 | 11.4.22 | −1.88 | 11.27.22 11:00 | −0.24 | 11.4.22 11:00 |
| | | Dec 2022 | 744 | −2.43 | 1.37 | −4.28 | 12.27.22 | −0.20 | 12.4.22 | −4.29 | 12.27.22 13:00 | −0.19 | 12.4.22 9:00 |
| | | Jan 2023 | 744 | −3.14 | 0.12 | −3.42 | 1.31.23 | −2.96 | 1.7.23 | −3.47 | 1.1.23 0:00 | −2.95 | 1.7.23 14:00 |
| | | Feb 2023 | 672 | −3.81 | 0.18 | −4.08 | 2.28.23 | −3.46 | 2.1.23 | −4.08 | 2.27.23 11:00 | −3.45 | 2.1.23 0:00 |
| | | Mar 2023 | 744 | −4.63 | 0.45 | −5.46 | 3.31.23 | −4.08 | 3.2.23 | −5.48 | 3.31.23 23:00 | −4.07 | 3.1.23 3:00 |
| | | Apr 2023 | 720 | −5.45 | 0.18 | −5.64 | 4.9.23 | −5.18 | 4.30.23 | −5.65 | 4.9.23 2:00 | −5.17 | 4.30.23 18:00 |
| | | May 2023 | 744 | −4.74 | 0.54 | −5.17 | 5.1.23 | −3.47 | 5.31.23 | −5.17 | 5.1.23 0:00 | −3.44 | 5.31.23 22:00 |
| | | Jun 2023 | 720 | −1.21 | 1.36 | −3.41 | 6.1.23 | −0.02 | 6.20.23 | −3.44 | 6.1.23 0:00 | −0.01 | 6.19.23 17:00 |
| | | Jul 2023 | 744 | 4.33 | 4.17 | −0.05 | 7.9.23 | 10.59 | 7.20.23 | −0.06 | 7.9.23 4:00 | 13.54 | 7.20.23 14:00 |
| | | Aug 2023 | 744 | 6.38 | 2.40 | 4.10 | 8.21.23 | 11.22 | 8.5.23 | 2.11 | 8.17.23 1:00 | 13.05 | 8.3.23 10:00 |
| | | Sep 2023 | 720 | −1.34 | 3.11 | −7.24 | 9.27.23 | 3.99 | 9.4.23 | −7.83 | 9.27.23 8:00 | 5.13 | 9.3.23 14:00 |
| | | Oct 2023 | 564 | −2.31 | 0.88 | −3.53 | 10.10.23 | −1.02 | 10.21.23 | −3.81 | 10.3.23 18:00 | −1.00 | 10.22.23 0:00 |

| | **Site** | **Month** | **n** | **Month mean** | | **Day mean minimum** | | **Day mean maximum** | | **Minimum measured** | | **Maximum measured** | |
|---|---|---|---|---|---|---|---|---|---|---|---|---|---|
| | | | | **Mean** | **s.d.** | **Min** | **Day** | **Max** | **Day** | **Min** | **Day, Hour** | **Max** | **Day, Hour** |
| **PAR ($\mu$mol m$^{-2}$ s$^{-1}$)** | Site 1 | Mar 2023 | 744 | 84.65 | 110.07 | 40.49 | 3.22.23 | 106.99 | 3.31.23 | 0.00 | 3.22.23 22:00 | 493.12 | 3.28.23 12:00 |
| | | Apr 2023 | 720 | 168.78 | 199.70 | 34.00 | 4.2.23 | 484.02 | 4.29.23 | 0.00 | 4.1.23 1:00 | 1699.11 | 4.29.23 15:00 |
| | | May 2023 | 744 | 73.69 | 175.33 | 0.69 | 5.7.23 | 507.22 | 5.27.23 | 0.00 | 5.10.23 22:00 | 1159.89 | 5.26.23 14:00 |
| | | Jun 2023 | 720 | 110.46 | 142.42 | 51.74 | 6.2.23 | 478.95 | 6.21.23 | 7.44 | 6.12.23 0:00 | 1273.53 | 6.21.23 19:00 |
| | | Jul 2023 | 744 | 356.09 | 326.12 | 114.84 | 7.1.23 | 637.58 | 7.3.23 | 21.93 | 7.18.23 23:00 | 1512.09 | 7.3.23 17:00 |
| | | Aug 2023 | 744 | 139.82 | 147.23 | 46.09 | 8.30.23 | 377.35 | 8.8.23 | 0.40 | 8.31.23 0:00 | 786.09 | 8.2.23 12:00 |
| | | Sep 2023 | 720 | 57.91 | 82.23 | 7.26 | 9.29.23 | 110.36 | 9.1.23 | 0.00 | 9.6.23 0:00 | 498.80 | 9.22.23 13:00 |
| | | Oct 2023 | 543 | 9.76 | 18.84 | 0.41 | 10.19.23 | 24.77 | 10.1.23 | 0.00 | 10.1.23 1:00 | 106.75 | 10.2.23 13:00 |
| | Site 2 | Mar 2023 | 744 | 115.81 | 131.08 | 97.44 | 3.29.23 | 143.25 | 3.26.23 | 0.00 | 3.27.23 1:00 | 461.73 | 3.30.23 12:00 |
| | | Apr 2023 | 720 | 220.74 | 205.43 | 99.08 | 4.2.23 | 340.88 | 4.29.23 | 0.00 | 4.1.23 1:00 | 896.90 | 4.23.23 14:00 |
| | | May 2023 | 744 | 378.06 | 256.39 | 168.74 | 5.5.23 | 548.61 | 5.27.23 | 35.23 | 5.5.23 0:00 | 1315.89 | 5.22.23 14:00 |
| | | Jun 2023 | 720 | 424.06 | 274.51 | 171.89 | 6.25.23 | 630.54 | 6.3.23 | 24.21 | 6.12.23 0:00 | 1406.97 | 6.16.23 12:00 |
| | | Jul 2023 | 744 | 405.67 | 268.72 | 154.26 | 7.28.23 | 621.16 | 7.3.23 | 28.58 | 7.29.23 0:00 | 1196.56 | 7.11.23 10:00 |
| | | Aug 2023 | 744 | 150.97 | 153.25 | 55.48 | 8.30.23 | 363.35 | 8.8.23 | 0.39 | 8.31.23 0:00 | 928.93 | 8.19.23 13:00 |
| | | Sep 2023 | 720 | 70.54 | 90.25 | 31.96 | 9.28.23 | 139.15 | 9.1.23 | 0.00 | 9.6.23 23:00 | 484.69 | 9.1.23 13:00 |
| | | Oct 2023 | 563 | 14.45 | 28.33 | 0.84 | 10.24.23 | 33.40 | 10.2.23 | 0.00 | 10.1.23 0:00 | 150.78 | 10.2.23 12:00 |
| | Site 3 | Jul 2023 | 744 | 352.18 | 229.59 | 135.21 | 7.28.23 | 570.25 | 7.14.23 | 27.28 | 7.19.23 1:00 | 1013.85 | 7.13.23 12:00 |
| | | Aug 2023 | 744 | 154.75 | 154.43 | 53.54 | 8.30.23 | 321.40 | 8.9.23 | 0.36 | 8.31.23 0:00 | 825.80 | 8.26.23 11:00 |
| | | Sep 2023 | 720 | 65.81 | 82.39 | 32.80 | 9.28.23 | 143.22 | 9.1.23 | 0.00 | 9.8.23 23:00 | 407.10 | 9.1.23 12:00 |
| | | Oct 2023 | 564 | 14.51 | 28.51 | 1.74 | 10.24.23 | 34.36 | 10.1.23 | 0.00 | 10.1.23 1:00 | 146.44 | 10.2.23 12:00 |

| | **Site** | **Month** | **n** | **Month mean** | | **Day mean minimum** | | **Day mean maximum** | | **Minimum measured** | | **Maximum measured** | |
|---|---|---|---|---|---|---|---|---|---|---|---|---|---|
| | | | | **Mean** | **s.d.** | **Min** | **Day** | **Max** | **Day** | **Min** | **Day, Hour** | **Max** | **Day, Hour** |
| **Humidity** | Site 1 | Mar 2023 | 744 | 75.55 | 9.54 | 63.80 | 3.25.23 | 88.76 | 3.30.23 | 55.86 | 3.23.23 14:00 | 95.11 | 3.30.23 19:00 |

| | | | | | | | | | | | | | |
|---|---|---|---|---|---|---|---|---|---|---|---|---|---|
| **(%)** | | Apr 2023 | 720 | 91.67 | 9.34 | 72.68 | 4.21.23 | 100.00 | 4.6.23 | 62.27 | 4.23.23 11:00 | 100.00 | 4.2.23 9:00 |
| | | May 2023 | 744 | 96.67 | 8.27 | 71.29 | 5.28.23 | 100.00 | 5.1.23 | 58.86 | 5.27.23 16:00 | 100.00 | 5.1.23 0:00 |
| | | Jun 2023 | 720 | 98.70 | 3.09 | 90.15 | 6.2.23 | 100.00 | 6.7.23 | 78.24 | 6.2.23 8:00 | 100.00 | 6.1.23 0:00 |
| | | Jul 2023 | 744 | 78.92 | 14.80 | 51.81 | 7.15.23 | 100.00 | 7.1.23 | 39.66 | 7.15.23 11:00 | 100.00 | 7.1.23 0:00 |
| | | Aug 2023 | 744 | 93.19 | 8.22 | 77.82 | 8.1.23 | 100.00 | 8.5.23 | 54.36 | 8.3.23 14:00 | 100.00 | 8.4.23 7:00 |
| | | Sep 2023 | 720 | 93.68 | 5.78 | 84.55 | 9.2.23 | 100.00 | 9.29.23 | 73.33 | 9.11.23 13:00 | 100.00 | 9.3.23 7:00 |
| | | Oct 2023 | 543 | 88.14 | 7.23 | 78.13 | 10.17.23 | 100.00 | 10.3.23 | 67.50 | 10.5.23 1:00 | 100.00 | 10.1.23 0:00 |
| | Site 2 | Mar 2023 | 744 | 87.40 | 7.86 | 73.38 | 3.26.23 | 91.52 | 3.31.23 | 66.72 | 3.26.23 16:00 | 100.00 | 3.31.23 20:00 |
| | | Apr 2023 | 720 | 89.89 | 9.78 | 71.46 | 4.5.23 | 98.97 | 4.28.23 | 44.47 | 4.5.23 11:00 | 100.00 | 4.2.23 11:00 |
| | | May 2023 | 744 | 90.28 | 11.28 | 64.58 | 5.19.23 | 98.81 | 5.31.23 | 53.89 | 5.3.23 12:00 | 100.00 | 5.4.23 22:00 |
| | | Jun 2023 | 720 | 92.47 | 8.79 | 69.67 | 6.2.23 | 100.00 | 6.19.23 | 56.91 | 6.2.23 7:00 | 100.00 | 6.6.23 22:00 |
| | | Jul 2023 | 744 | 83.33 | 14.83 | 53.12 | 7.6.23 | 99.98 | 7.1.23 | 41.92 | 7.6.23 10:00 | 100.00 | 7.1.23 1:00 |
| | | Aug 2023 | 744 | 96.76 | 6.81 | 79.45 | 8.9.23 | 100.00 | 8.12.23 | 60.43 | 8.3.23 13:00 | 100.00 | 8.1.23 17:00 |
| | | Sep 2023 | 720 | 96.41 | 4.97 | 89.73 | 9.14.23 | 100.00 | 9.29.23 | 76.77 | 9.28.23 0:00 | 100.00 | 9.1.23 0:00 |
| | | Oct 2023 | 563 | 92.70 | 8.71 | 78.52 | 10.23.23 | 100.00 | 10.15.23 | 55.32 | 10.19.23 20:00 | 100.00 | 10.1.23 0:00 |
| | Site 3 | Aug 2022 | 576 | 87.88 | 7.80 | 78.35 | 8.12.22 | 98.20 | 8.14.22 | 65.73 | 8.12.22 13:00 | 98.40 | 8.8.22 0:00 |
| | | Sep 2022 | 720 | 88.97 | 7.34 | 73.03 | 9.19.22 | 98.28 | 9.8.22 | 66.55 | 9.19.22 19:00 | 98.40 | 9.8.22 14:00 |
| | | Oct 2022 | 744 | 86.64 | 8.68 | 72.03 | 10.26.22 | 97.95 | 10.7.22 | 60.35 | 10.15.22 17:00 | 98.20 | 10.7.22 12:00 |
| | | Nov 2022 | 720 | 87.65 | 7.28 | 79.46 | 11.28.22 | 96.48 | 11.3.22 | 53.59 | 11.27.22 23:00 | 97.80 | 11.3.22 8:00 |
| | | Dec 2022 | 744 | 81.14 | 9.12 | 66.46 | 12.14.22 | 93.43 | 12.3.22 | 47.28 | 12.7.22 8:00 | 97.00 | 12.3.22 15:00 |
| | | Jan 2023 | 744 | 88.15 | 7.29 | 72.78 | 1.7.23 | 95.40 | 1.20.23 | 63.25 | 1.21.23 22:00 | 97.10 | 1.22.23 23:00 |
| | | Feb 2023 | 672 | 84.58 | 8.97 | 59.54 | 2.4.23 | 96.05 | 2.24.23 | 41.06 | 2.4.23 0:00 | 96.40 | 2.24.23 16:00 |
| | | Mar 2023 | 744 | 78.76 | 9.81 | 56.41 | 3.15.23 | 91.56 | 3.2.23 | 50.80 | 3.14.23 20:00 | 95.50 | 3.2.23 12:00 |
| | | Apr 2023 | 720 | 82.88 | 11.18 | 63.41 | 4.5.23 | 94.96 | 4.28.23 | 30.21 | 4.5.23 8:00 | 97.10 | 4.28.23 19:00 |
| | | May 2023 | 744 | 83.15 | 12.56 | 56.62 | 5.19.23 | 96.13 | 5.25.23 | 43.28 | 5.3.23 8:00 | 97.30 | 5.25.23 7:00 |
| | | Jun 2023 | 720 | 86.03 | 11.00 | 64.57 | 6.2.23 | 98.29 | 6.29.23 | 50.17 | 6.2.23 4:00 | 144.00 | 6.24.23 14:00 |
| | | Jul 2023 | 744 | 81.49 | 15.19 | 50.56 | 7.6.23 | 99.88 | 7.29.23 | 41.17 | 7.6.23 8:00 | 100.00 | 7.8.23 19:00 |
| | | Aug 2023 | 744 | 96.86 | 7.47 | 76.04 | 8.9.23 | 100.00 | 8.5.23 | 57.72 | 8.10.23 8:00 | 100.00 | 8.1.23 6:00 |
| | | Sep 2023 | 720 | 97.48 | 4.60 | 88.31 | 9.27.23 | 100.00 | 9.1.23 | 72.91 | 9.20.23 19:00 | 100.00 | 9.1.23 0:00 |
| | | Oct 2023 | 564 | 96.29 | 6.79 | 83.55 | 10.23.23 | 100.00 | 10.4.23 | 64.37 | 10.23.23 5:00 | 100.00 | 10.1.23 0:00 |

**Supplement S5.** The diurnal change of values of environmental parameters, maximum quantum yield ($F_V/F_M$; mean ± s.d.) and maximum possible relative electron transport rate ($rETR_{max}$; mean ± s.d.) during *in situ* measurement of photosynthetic activity. The presence of diurnal changes was tested by one-way ANOVA. Abbreviations: ANOVA – one-way ANOVA, n – number of cases, n.m. – not measured, PAR – photosynthetically active radiation, RH – relative air humidity, $T_{air}$ – air temperature, $T_{soil}$ – soil temperature. The statistically significant differences are marked in bold. Data used: averages per Petri dish and bowl.

| | Date, Time | n | $T_{air}$ (°C) | $T_{soil}$ (°C) | PAR (µmol m$^{-2}$ s$^{-1}$) | RH (%) | $F_V/F_M$ | $rETR_{max}$ |
|---|---|---|---|---|---|---|---|---|
| **2022 August** | | | | | | | | |
| Site 1 | 9/8, 17:00 | 2 | 10.4 | 12.1 | 364 | – | 0.273 ± 0.051 | 49.70 ± 9.28 |
| | 9/8, 23:00 | 6 | 6.1 | 7.9 | 36 | – | 0.520 ± 0.080 | 9.43 ± 1.45 |
| | 9/8, 5:00 | 6 | 8.0 | 8.2 | 119 | – | 0.525 ± 0.077 | 31.24 ± 4.59 |
| | 10/8, 11:00 | 6 | 10.3 | 10.9 | 231 | – | 0.368 ± 0.117 | 42.53 ± 13.53 |
| | 10/8, 17:00 | 6 | 9.1 | 10.7 | 175 | – | 0.287 ± 0.129 | 25.04 ± 11.30 |
| | ANOVA | | | | | | **P < 0.0001, F = 24.67** | **P < 0.0001, F = 51.85** |
| Site 2 | 9/8, 19:00 | 4 | 5.9 | 8.3 | 73 | – | 0.565 ± 0.073 | 20.74 ± 2.69 |
| | 10/8, 1:00 | 4 | 5.2 | 5.7 | 38 | – | 0.605 ± 0.061 | 11.62 ± 1.17 |
| | 10/8, 7:00 | 4 | 6.8 | 6.8 | 271 | – | 0.559 ± 0.057 | 75.84 ± 7.76 |
| | 10/8, 13:00 | 4 | 6.9 | 9.7 | 360 | – | 0.545 ± 0.061 | 97.96 ± 10.98 |
| | 10/8, 19:00 | 4 | 6.7 | 8.5 | 78 | – | 0.578 ± 0.059 | 22.46 ± 2.31 |
| | ANOVA | | | | | | P = 0.1545, F = 1.971 | **P < 0.0001, F = 596.3** |
| Site 3 | 9/8, 20:00 | 4 | 4.3 | 6.6 | 52 | 91.4 | 0.628 ± 0.047 | 16.46 ± 1.24 |
| | 10/8, 2:00 | 4 | 4.5 | 4.7 | 71 | 78.0 | 0.618 ± 0.067 | 21.93 ± 2.37 |
| | 10/8, 8:00 | 4 | 5.6 | 7.0 | 527 | 88.7 | 0.529 ± 0.073 | 139.51 ± 19.12 |
| | 10/8, 14:00 | 4 | 4.8 | 9.2 | 258 | 97.6 | 0.544 ± 0.052 | 70.16 ± 6.70 |
| | 10/8, 20:00 | 4 | 4.3 | 6.8 | 46 | 97.0 | 0.592 ± 0.072 | 13.55 ± 1.66 |
| | ANOVA | | | | | | **P < 0.0001, F = 12.90** | **P < 0.0001, F = 1211** |
| **2022 October** | | | | | | | | |
| Site 1 | 4/10, 8:00 | 6 | 1.9 | 1.1 | 0.1 | – | 0.597 ± 0.065 | 0.03 ± 0.00 |
| | 4/10, 12:00 | 6 | 1.8 | 1.1 | 86 | – | 0.496 ± 0.067 | 21.35 ± 2.90 |
| | 4/10, 17:00 | 6 | −0.1 | −0.1 | 22 | – | 0.581 ± 0.064 | 6.41 ± 0.71 |
| | ANOVA | | | | | | **P = 0.0002, F = 16.18** | **P < 0.0001, F = 1242** |
| **2023 August** | | | | | | | | |
| Site 1 | 5/8, 12:00 | 6 | 10.7 | 11.4 | 220 | 100.0 | 0.486 ± 0.092 | 53.56 ± 10.15 |
| | 5/8, 18:00 | 6 | 9.7 | 10.8 | 133 | 100.0 | 0.508 ± 0.077 | 33.64 ± 5.09 |
| | 6/8, 0:00 | 6 | 9.2 | 9.6 | 12 | 100.0 | 0.566 ± 0.068 | 3.47 ± 0.42 |
| | 6/8, 5:00 | 6 | 9.3 | 9.5 | 92 | 100.0 | 0.544 ± 0.074 | 25.04 ± 3.41 |
| | 6/8, 12:00 | 6 | 11.1 | 10.2 | 97 | 88.9 | 0.506 ± 0.072 | 24.59 ± 3.49 |
| | ANOVA | | | | | | **P = 0.0420, F = 2.907** | **P < 0.0001, F = 217.8** |
| Site 2 | 5/8, 14:00 | 4 | 10.8 | 12.1 | 272 | 100.0 | 0.524 ± 0.070 | 71.35 ± 9.56 |
| | 5/8, 19:00 | 4 | 10.0 | 11.2 | 62 | 100.0 | 0.570 ± 0.096 | 17.80 ± 3.01 |

| | | | | | | | | |
|---|---|---|---|---|---|---|---|---|
| | 6/8, 1:00 | 4 | 8.8 | 9.4 | 16 | 100.0 | 0.607 ± 0.084 | 4.85 ± 0.67 |
| | 6/8, 7:00 | 4 | 7.8 | 9.1 | 246 | 100.0 | 0.536 ± 0.080 | 65.96 ± 9.89 |
| | 6/8, 13:00 | 4 | 7.6 | 8.3 | 196 | 100.0 | 0.536 ± 0.066 | 52.49 ± 6.50 |
| | ANOVA | | | | | | P = 0.2640, F = 1.458 | **P < 0.0001, F = 218.2** |
| Site 3 | 5/8, 15:00 | 4 | 10.9 | 12.7 | 196 | 100.0 | 0.557 ± 0.054 | 54.44 ± 5.26 |
| | 5/8, 20:00 | 4 | 10.2 | 10.6 | 74 | 100.0 | 0.595 ± 0.069 | 21.91 ± 2.55 |
| | 6/8, 2:00 | 4 | 9.0 | 9.1 | 32 | 100.0 | 0.631 ± 0.058 | 10.22 ± 0.93 |
| | 6/8, 8:00 | 4 | 7.5 | 8.6 | 114 | 100.0 | 0.555 ± 0.054 | 31.56 ± 3.06 |
| | 6/8, 14:00 | 4 | 7.1 | 8.2 | 301 | 100.0 | 0.470 ± 0.067 | 70.67 ± 10.05 |
| | ANOVA | | | | | | **P < 0.0001, F = 24.46** | **P < 0.0001, F = 591.4** |
| **2023 October** | | | | | | | | |
| Site 1 | 23/10, 9:00 | 5 | −8.1 | −9.8 | 3.6 | 75.4 | 0.188 ± 0.035 | 0.34 ± 0.06 |
| | 23/10, 11:00 | 5 | −6.2 | −8.8 | 16 | 86.5 | 0.241 ± 0.053 | 1.99 ± 0.44 |
| | 23/10, 13:00 | 3 | −5.0 | −7.7 | 8.2 | 83.9 | 0.436 ± 0.257 | 1.80 ± 1.06 |
| | 23/10, 14:00 | 3 | −5.5 | −7.5 | 3.1 | 88.4 | 0.235 ± 0.046 | 0.37 ± 0.07 |
| | ANOVA | | | | | | **P = 0.0001, F = 17.69** | **P < 0.0001, F = 41.41** |

**Supplement S6.** The diurnal changes of the photosynthetic ($F_V/F_M$ and $rETR_{max}$; mean ± s.d., for n refer to Supplement S5) and environmental parameters (air and soil temperature, $T_{air}$, $T_{soil}$; photosynthetically active radiation, PAR; relative humidity, RH) at all the experimental sites in the studied periods in August 2022 and 2023.

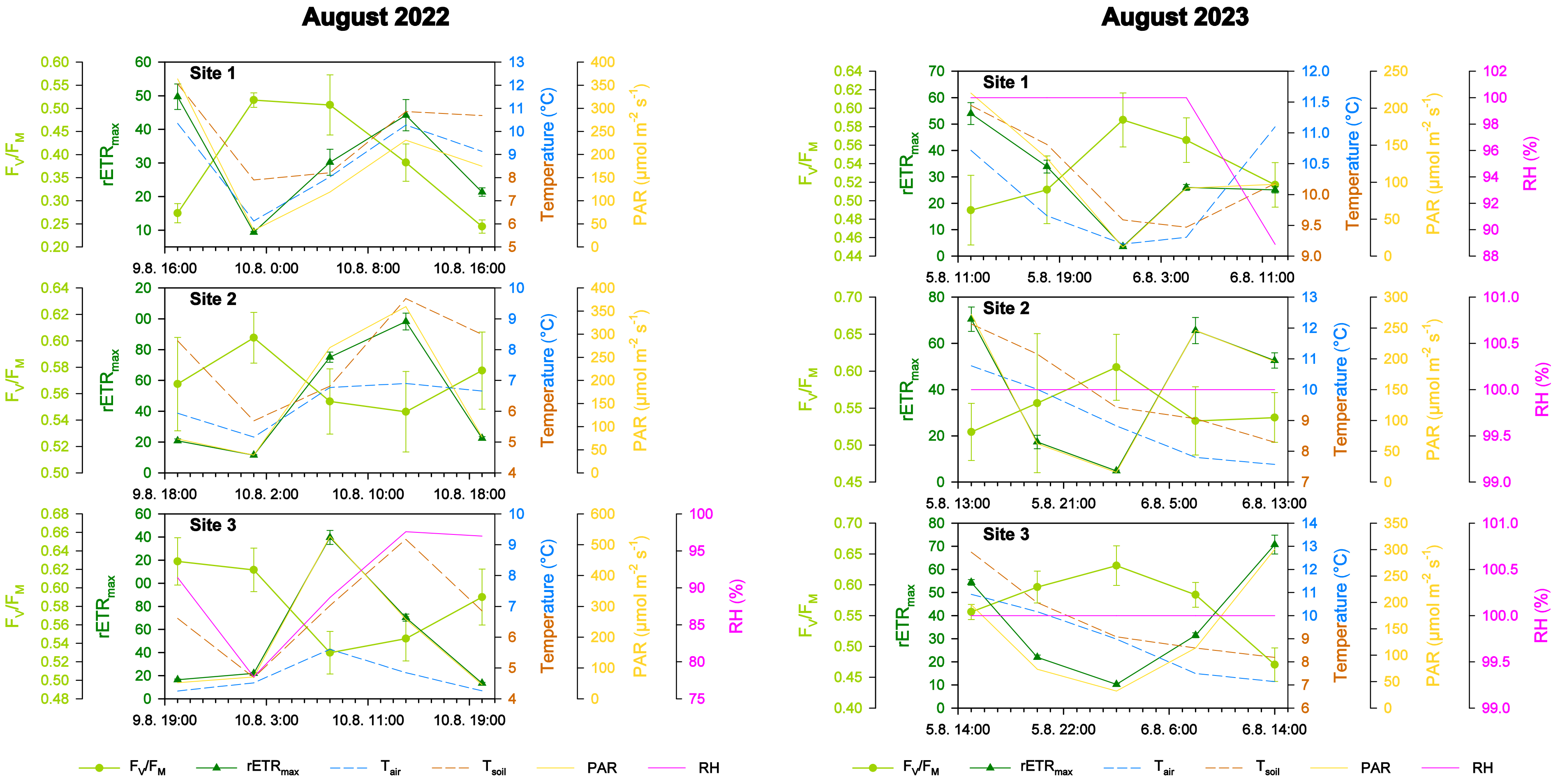

**Supplement S7**. Correlations of the $F_V/F_M$ and $rETR_{max}$ measured during diurnal cycles study with environmental data for the summer and autumn seasons 2022 and 2023. The statistically significant correlations are marked in bold.

| | | Air temperature | Soil temperature | Relative humidity | Photosynthetically active radiation |
|---|---|---|---|---|---|
| **$F_V/F_M$** | | | | | |
| **2022** | | | | | |
| Aug | Site 1 | P = 0.1308, r = −0.7662 | **P = 0.0248, r = −0.9241** | – | P = 0.2455, r = −0.7545 |
| | Site 2 | P = 0.0631, r = 0.8576 | P = 0.2056, r = 0.6810 | – | P = 0.0592, r = 0.8636 |
| | Site 3 | P = 0.0703, r = 0.8468 | P = 0.2173, r = 0.6686 | P = 0.5175, r = −0.3890 | **P = 0.0483, r = −0.8811** |
| Oct | Site 1 | P = 0.8100, r = 0.2940 | P = 0.7634, r = 0.3632 | – | P = 0.0659, r = −0.9946 |
| **2023** | | | | | |
| Aug | Site 1 | P = 0.1291, r = −0.7683 | **P = 0.0277, r = −0.9183** | P = 0.7038, r = 0.2348 | **P = 0.0389, r = −0.8974** |
| | Site 2 | P = 0.8573, r = −0.1123 | P = 0.6807, r = −0.2535 | P = 0.7356, r = 0.2092 | **P = 0.0262, r = −0.9213** |
| | Site 3 | P = 0.3588, r = 0.5295 | P = 0.6764, r = 0.2570 | P = 0.0683, r = 0.8497 | **P = 0.0101, r = −0.9584** |
| Oct | Site 1 | P = 0.2719, r = 0.7281 | P = 0.4323, r = 0.5677 | P = 0.7710, r = 0.2290 | P = 0.8395, r = 0.1605 |
| **$rETR_{max}$** | | | | | |
| **2022** | | | | | |
| Aug | Site 1 | **P = 0.0416, r = 0.8925** | P = 0.8390, r = 0.1268 | – | **P = 0.0363, r = 0.9019** |
| | Site 2 | P = 0.1395, r = 0.7557 | P = 0.4238, r = 0.4705 | – | **P < 0.0001, r = 1.000** |
| | Site 3 | **P = 0.0013, r = 0.9896** | P = 0.5267, r = 0.3811 | P = 0.9513, r = 0.0383 | **P < 0.0001, r = 0.9999** |
| Oct | Site 1 | P = 0.9058, r = 0.1475 | P = 0.8591, r = 0.2195 | – | **P = 0.0299, r = 0.9989** |
| **2023** | | | | | |
| Aug | Site 1 | P = 0.3355, r = 0.5512 | P = 0.0605, r = 0.8616 | P = 0.8666, r = 0.1050 | **P = 0.0001, r = 0.9976** |
| | Site 2 | P = 0.9112, r = −0.06982 | P = 0.9108, r = 0.07015 | P = 0.7491, r = −0.1983 | **P < 0.0001, r = 0.9996** |
| | Site 3 | P = 0.6928, r = −0.2437 | P = 0.9209, r = 0.06217 | P = 0.1437, r = 0.7507 | **P = 0.0008, r = 0.9925** |
| Oct | Site 1 | P = 0.4991, r = 0.5009 | P = 0.8267, r = 0.1733 | P = 0.6501, r = 0.3499 | P = 0.1249, r = 0.8751 |

**Supplement S8**. The changes of effective quantum yield ($\Phi_{PSII}$; mean ± s.d., n = 12) during recovery of photosynthetic activity in winter. The statistically significant differences were tested using Repeated Measures Analysis of Variance (RM ANOVA; n = 12). The letter in upper case indicates homologous groups recognized by Tukey HSD test for P = 0.05.

| **Recovery time (min)** | **March 2023, Thawing 1** | **March 2023, Thawing 2** | **March 2024, Thawing 1** | **March 2024, Thawing 2** |
|---|---|---|---|---|
| 0 | $0.470 \pm 0.098^{a}$ | $0.425 \pm 0.115^{a}$ | $0.195 \pm 0.174^{a,b}$ | $0.384 \pm 0.113^{a}$ |
| 5 | $0.461 \pm 0.081^{a}$ | $0.438 \pm 0.091^{a,b}$ | $0.221 \pm 0.148^{a}$ | $0.399 \pm 0.083^{a}$ |
| 10 | $0.502 \pm 0.075^{b}$ | $0.467 \pm 0.083^{b,c}$ | $0.264 \pm 0.124^{a,b}$ | $0.423 \pm 0.090^{a,b}$ |
| 15 | $0.522 \pm 0.065^{b,c}$ | $0.485 \pm 0.083^{c,d}$ | $0.296 \pm 0.155^{a,b}$ | $0.479 \pm 0.089^{c}$ |
| 20 | $0.531 \pm 0.070^{b,c,d}$ | $0.492 \pm 0.082^{c,d,e}$ | $0.272 \pm 0.148^{a,b}$ | $0.463 \pm 0.068^{b,c}$ |
| 25 | $0.537 \pm 0.075^{c,d,e}$ | $0.506 \pm 0.077^{d,e}$ | $0.294 \pm 0.165^{a,b}$ | $0.477 \pm 0.083^{c}$ |
| 30 | $0.540 \pm 0.075^{c,d,e}$ | $0.506 \pm 0.081^{d,e}$ | $0.294 \pm 0.178^{a,b}$ | $0.487 \pm 0.075^{c}$ |
| 35 | $0.553 \pm 0.073^{d,e}$ | $0.509 \pm 0.078^{d,e}$ | $0.358 \pm 0.153^{a,b}$ | $0.494 \pm 0.079^{c}$ |
| 40 | $0.558 \pm 0.063^{d,e}$ | $0.519 \pm 0.075^{d,e}$ | $0.330 \pm 0.183^{a,b}$ | $0.497 \pm 0.073^{c}$ |
| 45 | $0.559 \pm 0.066^{d,e}$ | $0.520 \pm 0.067^{d,e}$ | $0.350 \pm 0.143^{b}$ | $0.488 \pm 0.060^{c}$ |
| 50 | $0.563 \pm 0.067^{e}$ | $0.528 \pm 0.071^{e}$ | $0.361 \pm 0.155^{a,b}$ | $0.489 \pm 0.066^{c}$ |
| 55 | $0.565 \pm 0.071^{e}$ | $0.527 \pm 0.068^{e}$ | $0.364 \pm 0.173^{b}$ | $0.493 \pm 0.094^{c}$ |
| 60 | $0.567 \pm 0.064^{e}$ | $0.526 \pm 0.070^{e}$ | $0.343 \pm 0.177^{b}$ | $0.502 \pm 0.075^{c}$ |
| RM ANOVA | **P < 0.001, F = 28.65** | **P < 0.001, F = 19.03** | **P = 0.001, F = 2.998** | **P < 0.001, F = 16.75** |

**Supplement S9**. Relative abundances of photosynthesis-related transcripts (expressed in percentage of FPKM, fragments per kilobase of transcript per million fragments sequenced, mean ± s.d.) per study site and sampling season.

| | **Site1** | | | | **Site 2** | | | **Site 3** | | |
|---|---|---|---|---|---|---|---|---|---|---|
| **Gene** | **Aug22** (n = 5) | **Oct22** (n = 2) | **Aug23** (n = 4) | **Mar23** (n = 4) | **Aug22** (n = 5) | **Oct22** (n = 1) | **Aug23** (n = 4) | **Aug22** (n = 5) | **Oct22** (n = 2) | **Aug23** (n = 4) |
| PSII core | | | | | | | | | | |
| *PsbA* (D1 protein) | 64.016 ± 3.274 | 62.752 ± 3.270 | 75.079 ± 4.264 | 88.102 ± 3.686 | 67.291 ± 2.819 | 68.797 ± 0.000 | 76.008 ± 3.338 | 61.949 ± 2.297 | 74.007 ± 2.495 | 74.401 ± 2.747 |
| *PsbB* (CP47 protein) | 5.035 ± 0.976 | 4.517 ± 0.059 | 2.603 ± 0.262 | 1.118 ± 0.362 | 5.496 ± 0.652 | 4.507 ± 0.000 | 2.630 ± 0.324 | 5.214 ± 1.044 | 3.853 ± 0.699 | 3.058 ± 0.617 |
| *PsbC* (CP43 protein) | 10.675 ± 1.152 | 9.014 ± 1.668 | 4.707 ± 0.890 | 2.576 ± 0.689 | 7.646 ± 0.511 | 5.602 ± 0.000 | 4.295 ± 0.803 | 9.697 ± 1.411 | 7.547 ± 0.916 | 4.406 ± 0.505 |
| *PsbD* (D2 protein) | 5.220 ± 1.230 | 5.642 ± 0.631 | 2.910 ± 0.885 | 3.458 ± 1.935 | 5.137 ± 1.058 | 6.224 ± 0.000 | 3.408 ± 0.890 | 7.195 ± 1.847 | 5.438 ± 0.578 | 3.760 ± 1.208 |
| PSII small subunits | | | | | | | | | | |
| *PsbH* | 0.278 ± 0.085 | 0.285 ± 0.010 | 0.168 ± 0.031 | 0.064 ± 0.034 | 0.380 ± 0.161 | 0.333 ± 0.000 | 0.210 ± 0.058 | 0.495 ± 0.020 | 0.377 ± 0.120 | 0.333 ± 0.077 |
| *PsbI* | 0.185 ± 0.094 | 0.255 ± 0.131 | 0.117 ± 0.031 | 0.091 ± 0.069 | 0.102 ± 0.034 | 0.066 ± 0.000 | 0.085 ± 0.047 | 0.098 ± 0.031 | 0.042 ± 0.011 | 0.055 ± 0.033 |
| *PsbJ* | 0.073 ± 0.048 | 0.023 ± 0.008 | 0.025 ± 0.003 | 0.012 ± 0.008 | 0.064 ± 0.048 | 0.058 ± 0.000 | 0.023 ± 0.014 | 0.039 ± 0.007 | 0.053 ± 0.031 | 0.029 ± 0.013 |
| *PsbK* | 0.596 ± 0.305 | 0.403 ± 0.205 | 0.336 ± 0.112 | 0.137 ± 0.093 | 0.371 ± 0.107 | 0.166 ± 0.000 | 0.166 ± 0.055 | 0.295 ± 0.082 | 0.206 ± 0.125 | 0.318 ± 0.121 |
| *PsbL* | 0.069 ± 0.034 | 0.031 ± 0.000 | 0.042 ± 0.021 | 0.027 ± 0.026 | 0.056 ± 0.016 | 0.046 ± 0.000 | 0.030 ± 0.010 | 0.128 ± 0.043 | 0.075 ± 0.055 | 0.071 ± 0.018 |
| *PsbM* | 0.083 ± 0.065 | 0.011 ± 0.004 | 0.037 ± 0.023 | 0.018 ± 0.008 | 0.046 ± 0.032 | 0.015 ± 0.000 | 0.023 ± 0.007 | 0.036 ± 0.018 | 0.006 ± 0.006 | 0.018 ± 0.004 |
| *PsbR* | 0.140 ± 0.040 | 0.045 ± 0.021 | 0.042 ± 0.006 | 0.035 ± 0.018 | 0.112 ± 0.033 | 0.043 ± 0.000 | 0.046 ± 0.008 | 0.106 ± 0.043 | 0.103 ± 0.058 | 0.077 ± 0.031 |
| *PsbS* | 0.089 ± 0.036 | 0.072 ± 0.006 | 0.012 ± 0.011 | 0.043 ± 0.006 | 0.033 ± 0.009 | 0.070 ± 0.000 | 0.027 ± 0.017 | 0.033 ± 0.013 | 0.082 ± 0.010 | 0.035 ± 0.020 |
| *PsbT* | 0.029 ± 0.011 | 0.045 ± 0.017 | 0.037 ± 0.025 | 0.006 ± 0.005 | 0.044 ± 0.021 | 0.019 ± 0.000 | 0.032 ± 0.019 | 0.115 ± 0.080 | 0.063 ± 0.014 | 0.023 ± 0.005 |
| *PsbW* | 0.173 ± 0.160 | 0.128 ± 0.019 | 0.064 ± 0.013 | 0.052 ± 0.015 | 0.066 ± 0.020 | 0.070 ± 0.000 | 0.076 ± 0.022 | 0.060 ± 0.024 | 0.156 ± 0.022 | 0.090 ± 0.039 |
| *PsbX* | 0.001 ± 0.002 | 0.000 ± 0.000 | 0.001 ± 0.001 | 0.003 ± 0.002 | 0.001 ± 0.001 | 0.000 ± 0.000 | 0.016 ± 0.021 | 0.002 ± 0.003 | 0.014 ± 0.009 | 0.001 ± 0.002 |
| *PsbY* | 0.036 ± 0.026 | 0.021 ± 0.014 | 0.021 ± 0.005 | 0.019 ± 0.020 | 0.024 ± 0.012 | 0.008 ± 0.000 | 0.011 ± 0.007 | 0.019 ± 0.011 | 0.021 ± 0.012 | 0.012 ± 0.007 |
| *PsbZ* | 0.158 ± 0.042 | 0.097 ± 0.035 | 0.142 ± 0.033 | 0.066 ± 0.035 | 0.188 ± 0.053 | 0.155 ± 0.000 | 0.101 ± 0.032 | 0.278 ± 0.077 | 0.204 ± 0.051 | 0.136 ± 0.017 |
| Oxygen evolving complex | | | | | | | | | | |
| *PsbO* | 0.231 ± 0.137 | 0.307 ± 0.036 | 0.174 ± 0.006 | 0.110 ± 0.115 | 0.160 ± 0.058 | 0.155 ± 0.000 | 0.122 ± 0.031 | 0.194 ± 0.082 | 0.086 ± 0.017 | 0.127 ± 0.032 |
| *PsbP* | 0.012 ± 0.010 | 0.125 ± 0.040 | 0.024 ± 0.040 | 0.002 ± 0.003 | 0.001 ± 0.001 | 0.004 ± 0.000 | 0.005 ± 0.002 | 0.000 ± 0.000 | 0.001 ± 0.001 | 0.001 ± 0.001 |
| *PsbQ* | 0.006 ± 0.010 | 0.117 ± 0.040 | 0.018 ± 0.030 | 0.000 ± 0.000 | 0.002 ± 0.002 | 0.008 ± 0.000 | 0.006 ± 0.006 | 0.001 ± 0.002 | 0.003 ± 0.003 | 0.013 ± 0.006 |
| *PsbU* | 0.052 ± 0.020 | 0.086 ± 0.003 | 0.055 ± 0.007 | 0.027 ± 0.018 | 0.146 ± 0.129 | 0.054 ± 0.000 | 0.036 ± 0.009 | 0.190 ± 0.098 | 0.058 ± 0.009 | 0.049 ± 0.020 |
| Light harvesting complex | | | | | | | | | | |
| *Lhcb4* | 0.004 ± 0.004 | 0.007 ± 0.000 | 0.001 ± 0.003 | 0.000 ± 0.000 | 0.003 ± 0.004 | 0.000 ± 0.000 | 0.000 ± 0.001 | 0.002 ± 0.003 | 0.000 ± 0.000 | 0.003 ± 0.003 |
| *LhcA* | 0.370 ± 0.191 | 0.159 ± 0.019 | 0.091 ± 0.017 | 0.021 ± 0.013 | 0.393 ± 0.093 | 0.132 ± 0.000 | 0.092 ± 0.015 | 0.212 ± 0.044 | 0.070 ± 0.030 | 0.115 ± 0.012 |
| *LhcB* | 0.325 ± 0.130 | 0.149 ± 0.006 | 0.114 ± 0.043 | 0.029 ± 0.006 | 0.300 ± 0.044 | 0.155 ± 0.000 | 0.137 ± 0.020 | 0.198 ± 0.042 | 0.133 ± 0.012 | 0.175 ± 0.050 |
| PSII assembly | | | | | | | | | | |
| *Hcf136* | 0.005 ± 0.007 | 0.000 ± 0.000 | 0.000 ± 0.000 | 0.000 ± 0.000 | 0.001 ± 0.001 | 0.000 ± 0.000 | 0.000 ± 0.000 | 0.000 ± 0.001 | 0.009 ± 0.009 | 0.000 ± 0.001 |
| *Ohp1* | 0.001 ± 0.002 | 0.021 ± 0.006 | 0.002 ± 0.003 | 0.000 ± 0.000 | 0.000 ± 0.000 | 0.000 ± 0.000 | 0.000 ± 0.001 | 0.000 ± 0.000 | 0.000 ± 0.000 | 0.000 ± 0.000 |
| *Psb27* | 0.053 ± 0.027 | 0.052 ± 0.010 | 0.026 ± 0.007 | 0.013 ± 0.015 | 0.123 ± 0.176 | 0.039 ± 0.000 | 0.021 ± 0.007 | 0.039 ± 0.024 | 0.028 ± 0.013 | 0.031 ± 0.004 |
| *Psb28* | 0.000 ± 0.001 | 0.000 ± 0.000 | 0.000 ± 0.000 | 0.000 ± 0.000 | 0.001 ± 0.001 | 0.000 ± 0.000 | 0.001 ± 0.001 | 0.001 ± 0.001 | 0.000 ± 0.000 | 0.000 ± 0.000 |
| *Ycf48* | 0.006 ± 0.004 | 0.014 ± 0.014 | 0.005 ± 0.004 | 0.009 ± 0.005 | 0.026 ± 0.018 | 0.004 ± 0.000 | 0.013 ± 0.009 | 0.045 ± 0.033 | 0.021 ± 0.001 | 0.008 ± 0.005 |
| Carbon fixation | | | | | | | | | | |
| *RbcS* | 11.995 ± 2.512 | 15.524 ± 2.165 | 13.107 ± 4.043 | 3.935 ± 0.865 | 11.777 ± 1.282 | 13.191 ± 0.000 | 12.337 ± 2.817 | 13.335 ± 0.908 | 7.308 ± 1.288 | 12.605 ± 2.887 |
| Regulation/stress response | | | | | | | | | | |
| *Cor413pm1* | 0.014 ± 0.027 | 0.007 ± 0.003 | 0.002 ± 0.002 | 0.002 ± 0.003 | 0.005 ± 0.007 | 0.077 ± 0.000 | 0.033 ± 0.024 | 0.003 ± 0.003 | 0.029 ± 0.006 | 0.026 ± 0.028 |
| *Elip* | 0.072 ± 0.035 | 0.092 ± 0.011 | 0.038 ± 0.013 | 0.030 ± 0.014 | 0.008 ± 0.009 | 0.004 ± 0.000 | 0.007 ± 0.006 | 0.021 ± 0.013 | 0.008 ± 0.001 | 0.023 ± 0.013 |

**Supplement S10**. Results of two-factor ANOVA ($n_{(Aug22)} = 5$, $n_{(Oct22)} = 2$, $n_{(Mar23,\ Aug23)} = 4$) assessing the impact of site (Site 1 × Site 2 × Site 3) and sampling season (Aug22 × Oct22 × Mar23 × Aug23) on photosynthesis- and stress-related transcripts represented by FPKM numbers (fragments per kilobase of transcript per million fragments sequenced).

| Gene | Site | Sampling season | Interaction (Site × Season) |
|---|---|---|---|
| PSII core | | | |
| ***PsbA* (D1 protein)** | **P = 0.0132, F = 5.1390** | **P = 0.0013, F = 7.0600** | P = 0.6764, F = 0.5850 |
| ***PsbB* (Cp47 protein)** | **P = 0.0101, F = 5.5090** | **P < 0.0001, F = 14.1020** | P = 0.5549, F = 0.7690 |
| ***PsbC* (Cp43 protein)** | **P = 0.0104, F = 5.4760** | **P < 0.0001, F = 24.9630** | P = 0.3422, F = 1.1810 |
| ***PsbD* (D2 protein)** | **P = 0.0030, F = 7.3290** | **P < 0.0001, F = 13.5950** | P = 0.2054, F = 1.5950 |
| PSII small subunits | | | |
| ***PsbH*** | **P = 0.0004, F = 10.5500** | **P = 0.0001, F = 10.8400** | P = 0.2256, F = 1.5200 |
| ***PsbI*** | P = 0.8927, F = 0.1140 | **P = 0.0031, F = 5.9820** | P = 0.9309, F = 0.2090 |
| ***PsbJ*** | P = 0.0835, F = 2.7360 | **P < 0.0001, F = 11.9730** | P = 0.8416, F = 0.3500 |
| ***PsbK*** | P = 0.5570, F = 0.5990 | **P < 0.0001, F = 14.9600** | P = 0.7250, F = 0.5160 |
| ***PsbL*** | **P < 0.0001, F = 14.9080** | **P < 0.0001, F = 14.3910** | **P = 0.0400, F = 2.9280** |
| ***PsbM*** | P = 0.6476, F = 0.4420 | **P = 0.0055, F = 5.3010** | P = 0.9975, F = 0.0350 |
| ***PsbR*** | P = 0.1730, F = 1.8790 | **P = 0.0007, F = 7.8740** | P = 0.9972, F = 0.0370 |
| ***PsbS*** | P = 0.8692, F = 0.1410 | **P = 0.0022, F = 6.3660** | P = 0.2202, F = 1.5390 |
| ***PsbT*** | **P = 0.0428, F = 3.5650** | P = 0.0752, F = 2.5790 | P = 0.1656, F = 1.7680 |
| ***PsbW*** | P = 0.8354, F = 0.1810 | **P = 0.0177, F = 4.0300** | P = 0.5357, F = 0.8010 |
| *PsbX* | P = 0.2790, F = 1.3390 | P = 0.8000, F = 0.3360 | P = 0.1710, F = 1.7420 |
| ***PsbY*** | P = 0.7650, F = 0.2710 | **P = 0.0273, F = 3.5800** | P = 0.9724, F = 0.1240 |
| ***PsbZ*** | **P = 0.0011, F = 8.9750** | **P = 0.0001, F = 11.6170** | P = 0.0625, F = 2.5580 |
| Oxygen evolving complex | | | |
| ***PsbO*** | P = 0.7077, F = 0.3500 | **P = 0.0085, F = 4.8180** | P = 0.8740, F = 0.3020 |
| ***PsbP*** | **P = 0.0097, F = 5.5700** | **P = 0.0055, F = 5.3070** | **P = 0.0247, F = 3.3390** |
| ***PsbQ*** | P = 0.1847, F = 1.8040 | **P = 0.0035, F = 5.8310** | **P = 0.0033, F = 5.1980** |
| *PsbU* | P = 0.1428, F = 2.1000 | P = 0.0697, F = 2.6520 | P = 0.4592, F = 0.9350 |
| Light harvesting complex | | | |
| *Lhcb4* | P = 0.8810, F = 0.1270 | P = 0.2510, F = 1.4500 | P = 0.7510, F = 0.4790 |
| ***LhcA*** | **P = 0.0125, F = 5.2140** | **P < 0.0001, F = 17.1710** | P = 0.3211, F = 1.2330 |
| ***LhcB*** | **P = 0.0251, F = 4.2620** | **P < 0.0001, F = 13.9780** | P = 0.7730, F = 0.4480 |
| PSII assembly | | | |
| *Hcf136* | P = 0.9380, F = 0.0640 | P = 0.1930, F = 1.6950 | P = 0.2110, F = 1.5730 |
| ***Ohp1*** | **P = 0.0054, F = 6.4260** | **P = 0.0001, F = 11.2450** | **P < 0.0001, F = 10.1770** |
| *Psb27* | P = 0.3660, F = 1.0460 | P = 0.5040, F = 0.8020 | P = 0.7800, F = 0.4380 |
| *Psb28* | P = 0.0840, F = 2.7280 | P = 0.5130, F = 0.7860 | P = 0.8430, F = 0.3480 |
| ***Ycf48*** | **P = 0.0423, F = 3.5810** | P = 0.0548, F = 2.8860 | P = 0.2259, F = 1.5190 |
| Carbon fixation | | | |
| ***RbcS*** | **P = 0.0020, F = 7.9790** | **P < 0.0001, F = 11.6690** | P = 0.1432, F = 1.8850 |
| Regulation/stress response | | | |
| *Cor413pm1* | P = 0.3700, F = 1.0330 | P = 0.5350, F = 0.7450 | P = 0.1860, F = 1.6750 |
| ***Elip*** | **P = 0.0112, F = 5.3620** | **P = 0.0056, F = 5.2870** | P = 0.5648, F = 0.7540 |

**Supplement S11**. RDA analyses showing correlation among relative abundances of photosynthesis-related transcripts (explained variables: relative abundances of photosynthesis-related genes; arrows) and environmental parameters (explaining variables: sampling season; red symbols) and separation of gene expression at individual sites. The total variation is 480 (Site 1) / 320 (Site 2) / 341 (Site 3), explanatory variables account for 51.28 % / 37.82 % / 41.73 % of explained variation. Monte Carlo Permutation test results: P = 0.002 / P = 0.004/ P = 0.008, pseudo-F = 1.9 / 1.7 / 1.7 (first axis); P = 0.002 / P = 0.022 / P = 0.002, pseudo-F = 1.9 / 1.7 / 1.7 (all axes).

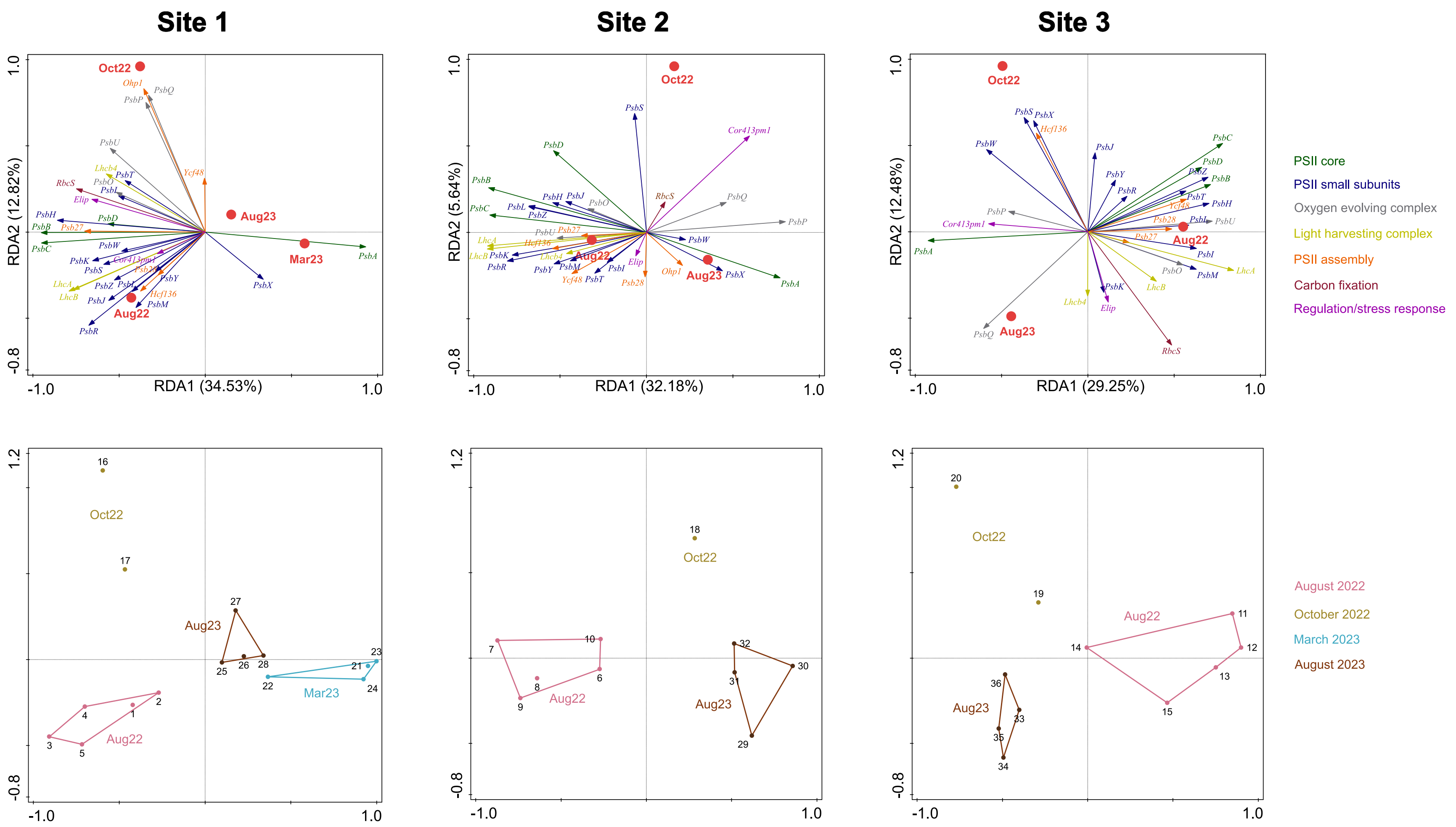
